## Supplementary Figures for "Conformational plasticity across phylogenetic clusters of RND multidrug efflux pumps and its impact on substrate specificity"

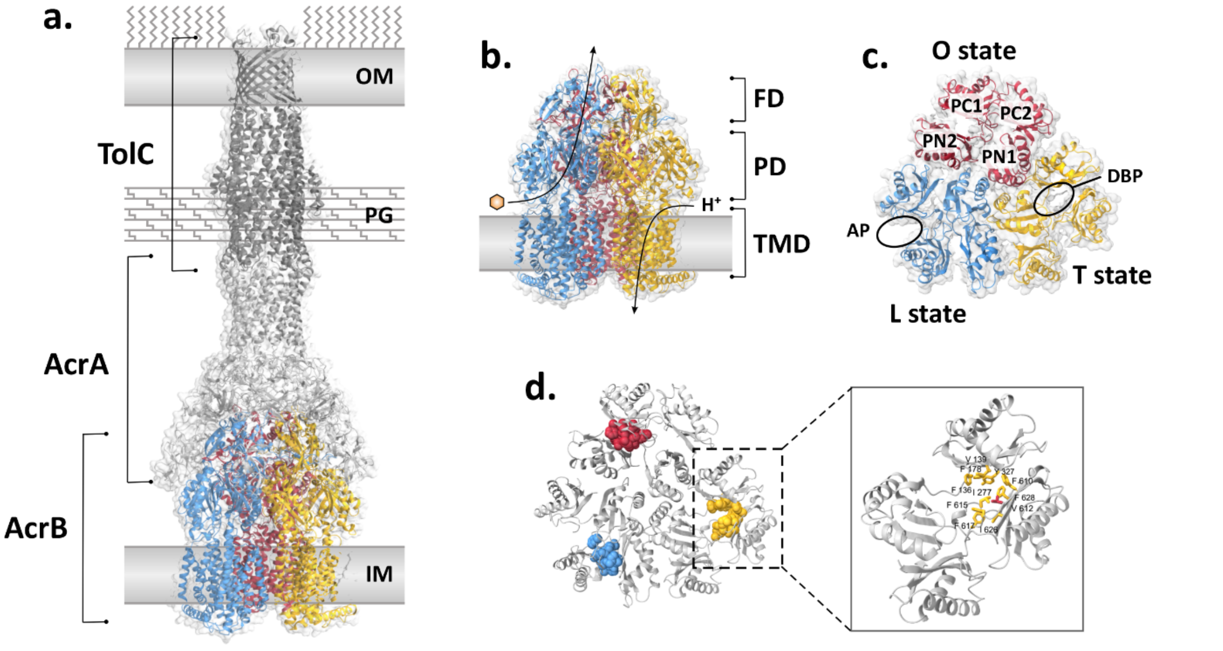


**Figure S1: The AcrAB-TolC system from *E. coli*.** (**a.**) AcrAB-TolC is a constitutively expressed tripartite multidrug efflux system from *E. coli*. It consists of the inner membrane Resistance Nodulation and cell Division (RND) transporter AcrB, the pore forming outer membrane factor TolC and the periplasmic adaptor AcrA. (**b.**) AcrB is a drug/proton antiporter that energises the drug export and the determinant of the substrate specificity of the entire tripartite system. It forms a homotrimer in the inner membrane and each monomer can adopt one of three distinct conformational states called loose (L), tight (T) and open (O). During the catalytic transport cycle every protomer cycles through these three states in a concerted fashion with the other neighbouring protomers (LTO -> TOL -> OLT -> LTO). AcrB can be subdivided into a transmembrane domain (TMD) that contains a proton translocation network, a porter domain (PD) that contains the substrate binding pockets, and a funnel domain (FD) from where drug substrates are entering into the AcrA-TolC channel. (**c.**) The porter domain can be divided into four subdomains (PN1, PN2, PC1 and PC2). During the transport cycle, these domains move as rigid body units relative to each other thus inducing the opening and closing of the substrate binding pockets. Substrates are bound in the access pocket (AP) in the L state and in the deep binding pocket (DBP) in the T state. In the O state these pockets are closed, but an exit channel leading to the AcrA-TolC funnel is opened. (**d.**) The DBP is lined by hydrophobic, mostly aromatic, residues that form an open pocket cleft in the T state. In the O and L states this pocket is closed, and a close packing of the DPB residues is observed. Colours in all subfigures: blue – L state, yellow – T state, red – O state.

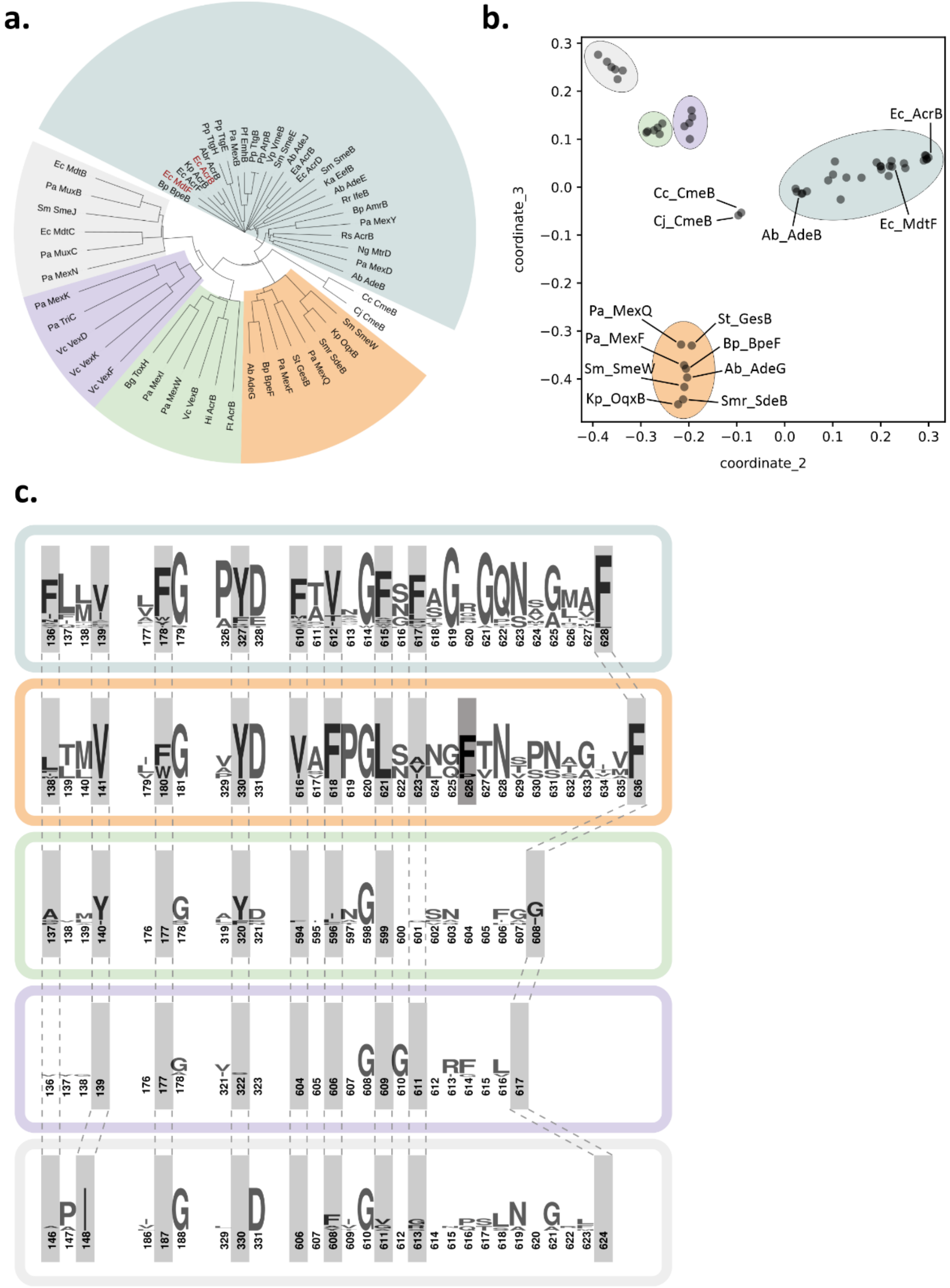


**Figure S2: Conservation of deep binding pocket (DBP) residues in RND proteins from Gram-negative bacteria.** A phylogenetic tree (**a.**) and a cc-analysis map of pairwise sequence alignment (**b.**) of the same set of HAE-1 RND amino acid sequences (table S1). With both methods the same five clusters were identified (highlighted by the same colour in all subfigures). The CmeB genes did not cluster with any of the other genes. The AcrB cluster (cyan) comprises several subclusters such as the MdtF and AdeB subclusters. (**c.**) Consensus sequence of the DBP residues for each cluster. The residues numbering corresponds to the sequences of following cluster members: AcrB (cyan), OqxB (orange), MexW (green), VexF (purple) and MdtB (grey). DBP residues are highlighted in grey and dashed lines connect the equivalent residues in each consensus sequence. The DBP residues are conserved in the AcrB and OqxB clusters (cyan and orange, respectively), but not in the remaining three clusters. The position equivalent to AcrB_F617 is not conserved in the OqxB cluster but is compensated by F626 (darker grey) in the latter. Abbreviations: Ec: *Escherichia coli*; Rr: *Rhizobium radiobacter*; Ab: *Acinetobacter baumannii*; Ng: *Neisseria gonorrhoeae*; Pa: *Pseudomonas aeruginosa*; Pp: *Pseudomonas putida*; Sm: *Stenotrophomonas maltophilia*; Ka: *Klebsiella aerogenes*; Bg: *Burkholderia glumae*; Cj: *Campylobacter jejuni*; Cc: *Campylobacter coli*; Bp: *Burkholderia pseudomallei*; St: *Salmonella typhimurium*; Vp: *Vibrio parahaemolyticus*; Ft: *Francisella tularensis*; Vc: *Vibrio cholerae*; Smr: *Serratia marcescens*; Rs: *Ralstonia solanacearum*; Abr: *Alcanivorax borkumensis*; Kp: *Klebsiella pneumoniae*; Ea: *Erwinia amylovora*; Hi: *Haemophilus influenzae*; Pf: *Pseudomonas fluorescens*

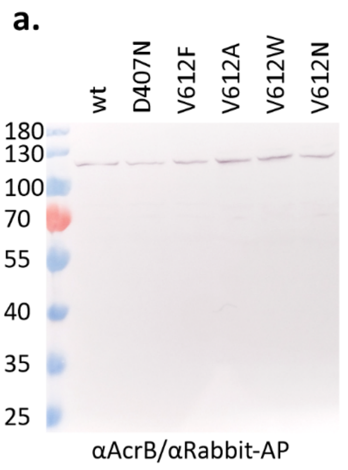


b.

| | wt | V612F | V612W | V612N | V612A | D407N |
|---|---|---|---|---|---|---|
| CAM | 1,00 | 2,00 | 1,67 | 1,67 | 1,67 | 0,34 |
| TIA | 1,00 | 2,00 | 1,67 | 2,00 | 1,00 | 0,50 |
| LIN | 1,00 | 1,80 | 2,00 | 1,33 | 1,83 | 0,06 |
| TET | 1,00 | 1,00 | 1,00 | 1,00 | 1,33 | 0,36 |
| OxTET | 1,00 | 2,00 | 2,00 | 2,00 | 2,00 | 0,50 |
| TIG | 1,00 | 0,82 | 1,09 | 1,00 | 1,09 | 0,55 |
| CITET | 1,00 | 1,00 | 1,00 | 1,00 | 1,00 | 0,25 |
| DXC | 1,00 | 1,00 | 1,00 | 1,00 | 1,00 | 0,15 |
| MIN | 1,00 | 0,50 | 0,72 | 0,89 | 0,72 | 0,31 |
| DIC | 1,00 | 0,50 | 0,50 | 0,50 | 1,00 | 0,07 |
| OXA | 1,00 | 0,50 | 0,67 | 0,58 | 0,83 | 0,02 |
| PIP | 1,00 | 0,31 | 0,67 | 0,58 | 0,83 | 0,22 |
| SDS* | | | | | | |
| ERY | 1,00 | 0,44 | 0,50 | 0,50 | 0,75 | 0,06 |
| CLA | 1,00 | 0,50 | 0,92 | 0,50 | 1,00 | 0,11 |
| NOV | 1,00 | 0,45 | 0,64 | 0,72 | 1,56 | 0,18 |
| FUA | 1,00 | 0,45 | 0,75 | 0,50 | 1,00 | 0,11 |
| H33342 | 1,00 | 0,34 | 0,38 | 0,83 | 0,75 | 0,20 |

MIC variant/MIC wt (colour scale: 2 1 0)

**Figure S3: Phenotype characterisation of AcrB_V612 variants.** For the functional characterisation of AcrB V612 variants, BW25113 Δ*acrB* cells were transformed with a pET24 vector encoding the AcrB variants. For each functional assay, Western blot samples were prepared from membranes isolated from the used clones to verify the presence of the AcrB variants. Western blot membranes were stained with a rabbit anti-AcrB primary antibody and an anti-rabbit-alkaline phosphatase conjugated secondary antibody [1]. A representative blot is shown in (**a**.). For all variants a band for the AcrB monomer (calculated molecular weight of 114 kDa) is visible. (**b.**) Minimal inhibitory concentration (MIC) assay. Bacterial cultures were exposed to a serial dilution of a toxic substrate. The inactive AcrB variant D407N [2] was used as negative control. The first dilution step at which no growth was detected was determined (MIC) and normalised to the wildtype (wt) MIC. Values higher than one (indicated with green colour) represent a higher MIC and values lower than one (purple) represent a lower MIC compared to the wt for each given toxin. The figure represents summary data of at least three biological replicates. The mean MIC values used for the figure generation are summarized in table S3 and the individual MIC values of all replicates are given in the source data. *For SDS, the MIC value of the wt exceeded the maximal tested concentration, therefore MIC values of the cells harbouring AcrB variants could not be normalised to the wt. Nonetheless, a clear reduction of the resistance against SDS was detected for cells harbouring the V612F and V612W variants (table S3). Abbreviations of the tested drugs, given at the left side of the figure, are specified in table S2.

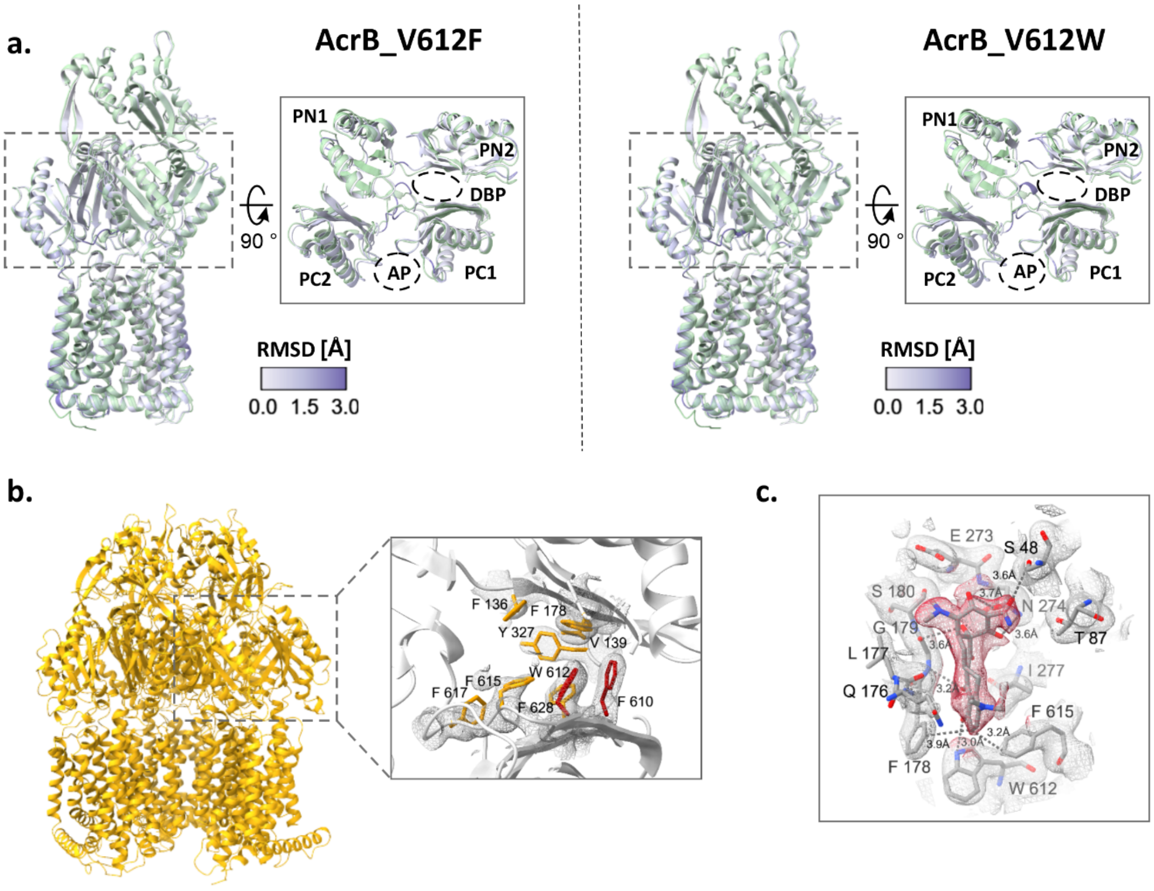


**Figure S4: Comparison of V612F/W structures with AcrB wildtype.** (**a.**) The V612F/W structures were aligned with the AcrB wildtype structure in the T state (PDB ID: 4dx5, green) and coloured by the RMSD values as indicated in the colour key. Inset: top view of the porter domain. The PN1, PN2, PC1 and PC2 subdomains are indicated. AP: Access Pocket, DBP: Deep Binding Pocket (**b.**) Crystallographic structure of AcrB V612W in the TTT state. The deep binding pocket residues in AcrB_V612W are shown in the inlet on the right. The crystallographic 2Fo-Fc map is shown at $\sigma 1$ (mesh). (**c.**) Minocycline interactions in the deep binding pocket of AcrB V612W. The crystallographic 2Fo-Fc map is shown at $\sigma 1$ (mesh) and the densities for minocycline (shown as sticks) are highlighted in red. Residues with at least one atom within 4 Å distance of minocycline are shown as sticks and indicated with single letter amino acid code and position number. The distances between the side chains and minocycline (dashed lines) are indicated. Carbon atoms are given in grey, oxygen in red, and nitrogen in blue.

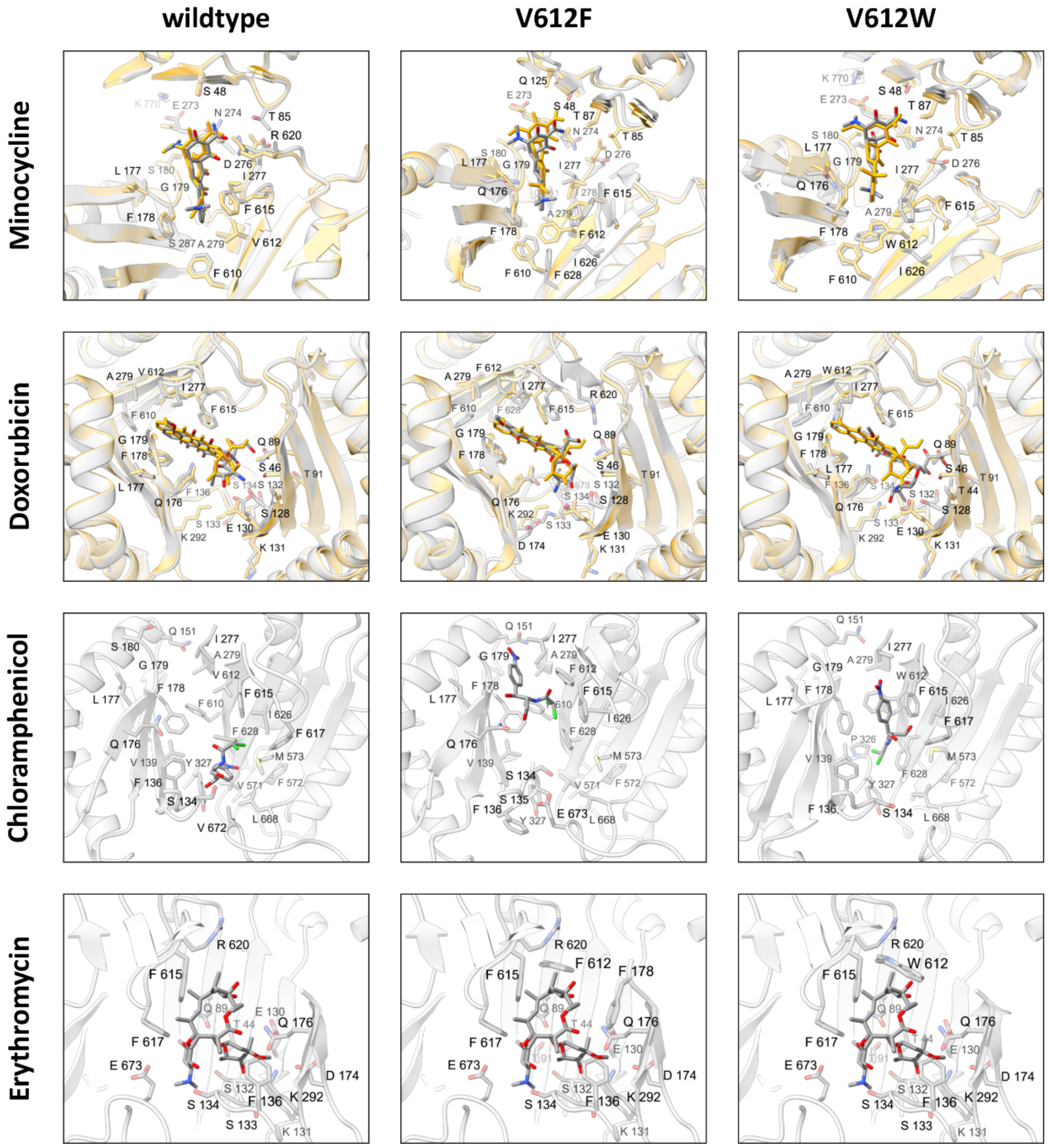


**Figure S5: Docking of AcrB substrates in the deep binding pocket of AcrB wildtype and the V612F/W variants.** The substrates and potential interacting residues (< 4 Å) are shown as sticks. All docking poses are coloured by atom type with carbon atoms in grey, oxygen in red, and nitrogen in blue. For minocycline, the docking poses were overlayed with the crystallographic structures of the wildtype (PDB ID: 4dx5) or the crystallographic structures of the respective V612 variant (this study, yellow). For doxorubicin, all docking poses were overlayed with the crystallographic structure of the AcrB wildtype (PDB ID: 4dx7, yellow). For a detailed discussion of the results see the supplementary information.

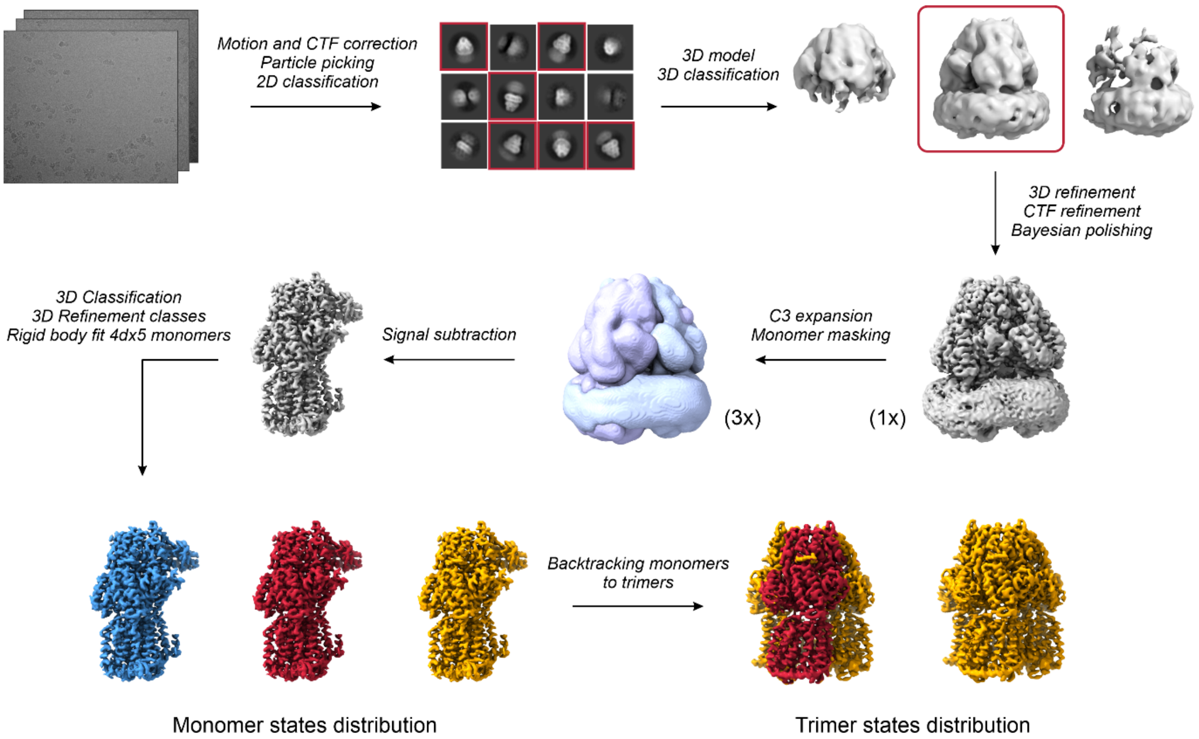


**Figure S6: Processing pipeline for cryo-EM datasets.** Micrographs from vitrified samples were recorded on a Titan Krios cryo-TEM operating at 300 kV. Beam-induced motion correction and CTF estimation were performed prior to template based automated particle picking. Several iterative rounds of 2D and 3D classification were performed to remove poor-quality particles, and a 3D volume of the trimer was reconstituted. Utilising the C3 pseudosymmetry through the central axis of the protein, a symmetry expansion was performed, and a soft monomer mask was used to subtract the signal of the two other monomers. Several 3D classification rounds were performed to obtain the maximum number of classes (minimum 3) with the best resolution. Class averages were refined. The conformational state of the refined 3D volumes was assessed by comparison with the AcrB states from the best resolved (1.9 Å) AcrB crystallographic structure (PDB ID: 4dx5). A custom MATLAB script was used to calculate the trimer composition based on the position of the monomer particles.

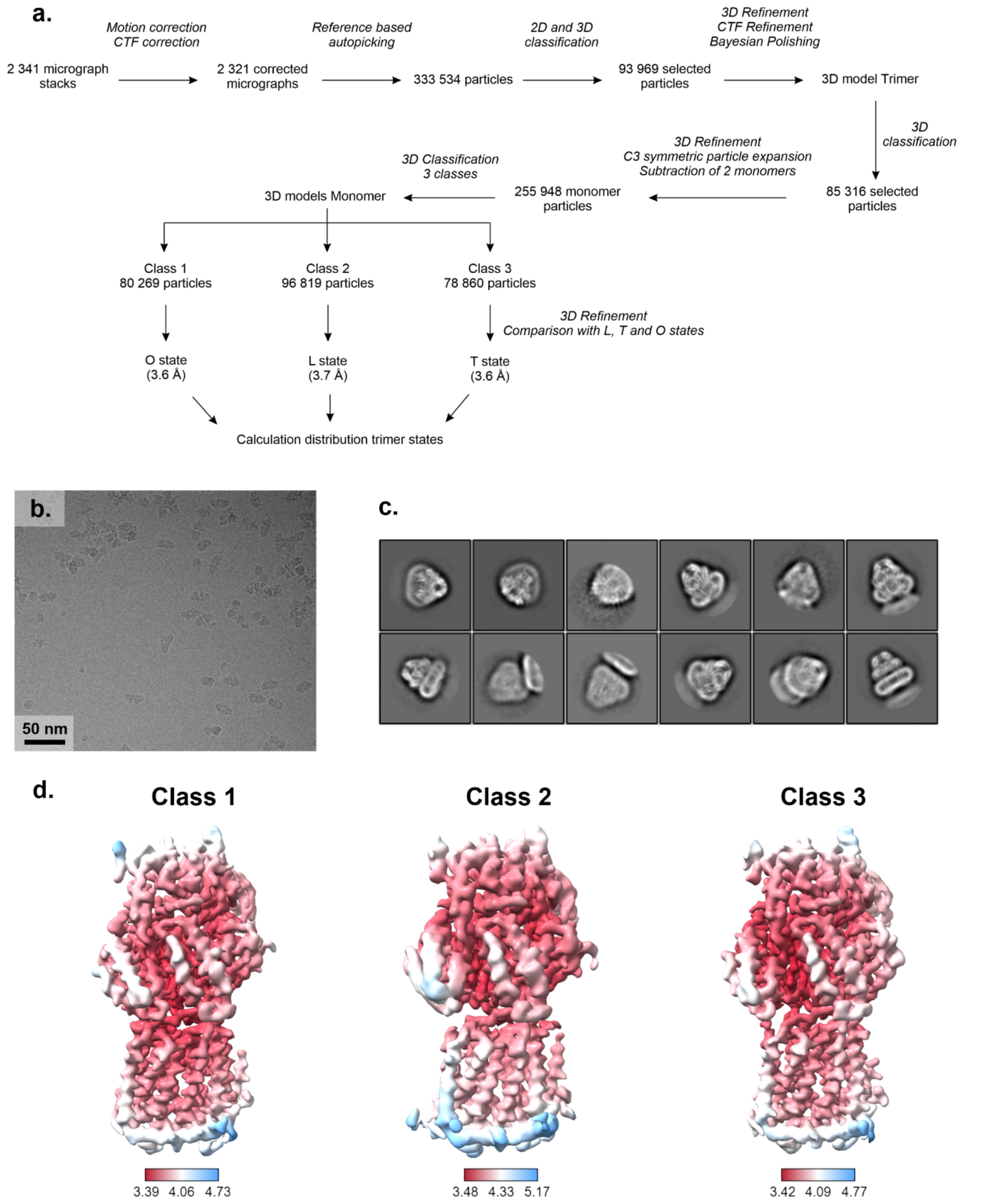


**Figure S7: Cryo-EM data of AcrB wildtype solubilised in DDM (a.)** Processing pipeline for evaluation of the distribution of conformational states. The processing was performed in Relion [3]. The conformational state of each class was determined by comparison with the three monomers of the best resolved asymmetric LTO structure of AcrB wildtype (PDB ID: 4dx5). Representative micrograph and 2D classes from the dataset are shown in (**b.**). and (**c.**), respectively. The local resolution of the 3D volumes of the monomer class averages is depicted in (**d.**)

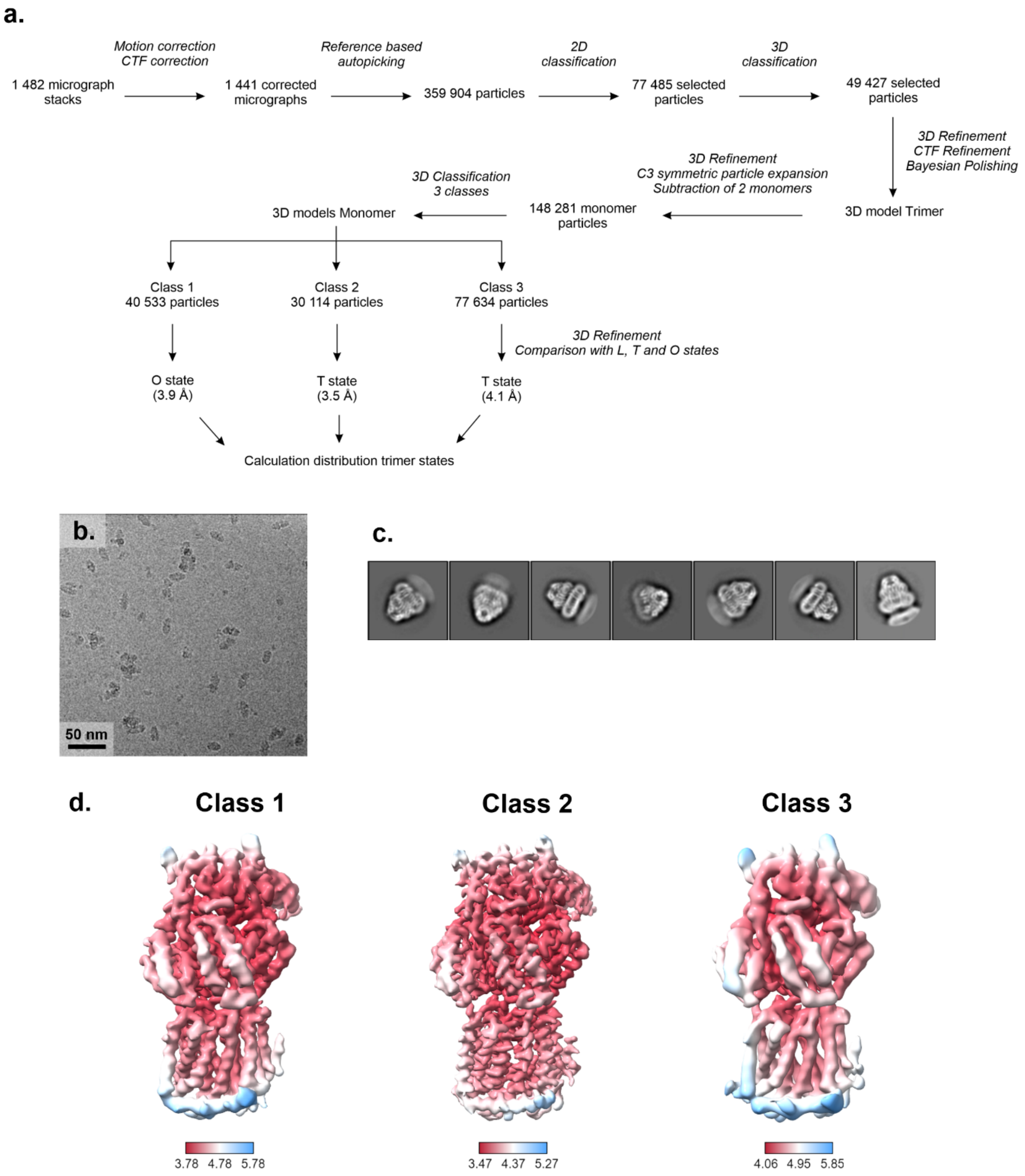


**Figure S8: Cryo-EM data of AcrB V612F solubilised in DDM. (a.)** Processing pipeline for evaluation of the distribution of conformational states. The processing was performed in Relion [3]. The conformational state of each class was determined by comparison with the three monomers of the best resolved asymmetric LTO structure of AcrB wildtype (PDB ID: 4dx5). Representative micrograph and 2D classes from the dataset are shown in (**b.**). and (**c.**), respectively. The local resolution of the 3D volumes of the monomer class averages is depicted in (**d.**)

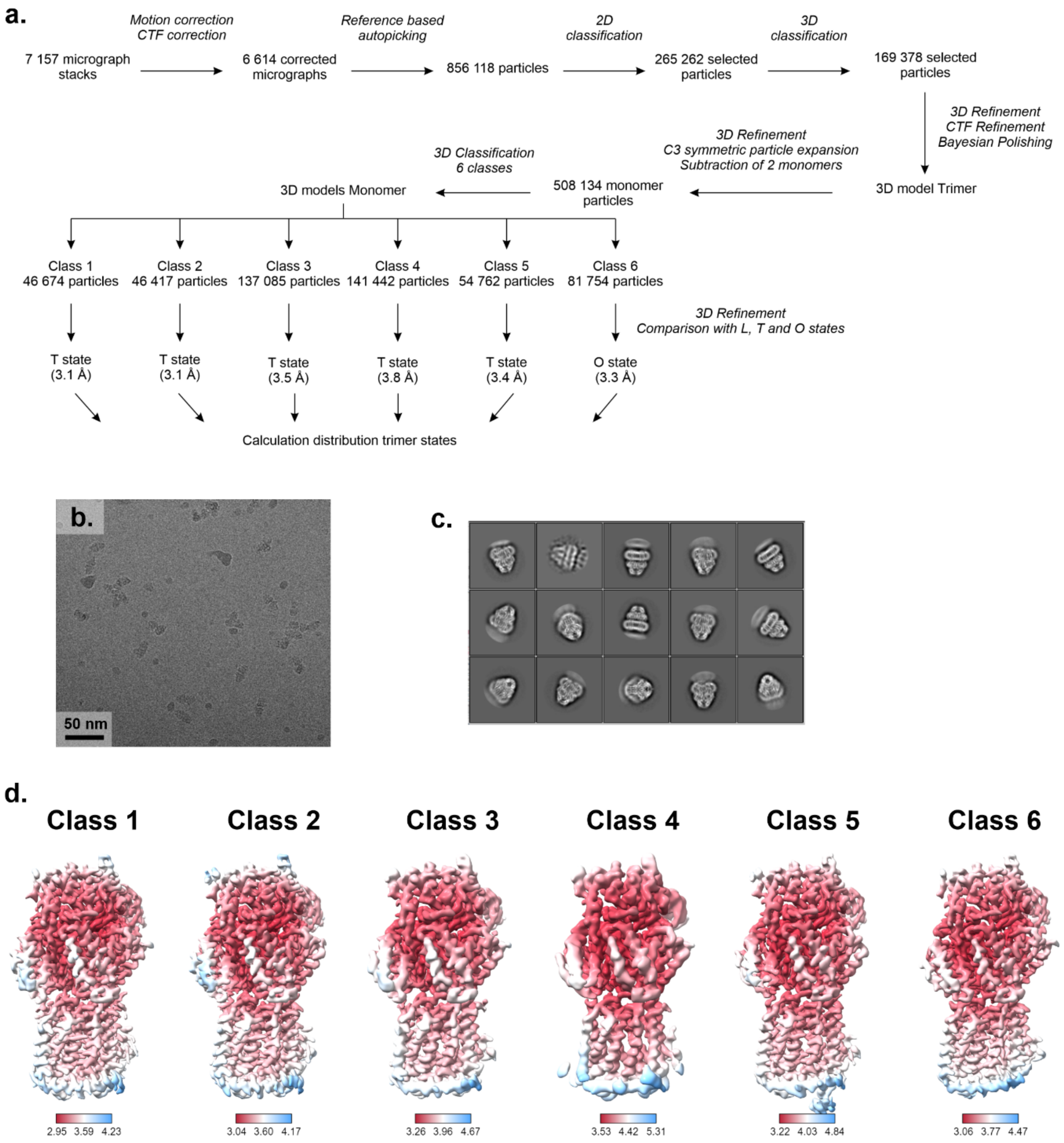


**Figure S9: Cryo-EM data of AcrB V612W solubilised in DDM. (a.**) Processing pipeline for evaluation of the distribution of conformational states. The processing was performed in Relion [3]. The conformational state of each class was determined by comparison with the three monomers of the best resolved asymmetric LTO structure of AcrB wildtype (PDB ID: 4dx5). Representative micrograph and 2D classes from the dataset are shown in (**b.**). and (**c.**), respectively. The local resolution of the 3D volumes of the monomer class averages is depicted in (**d.**)

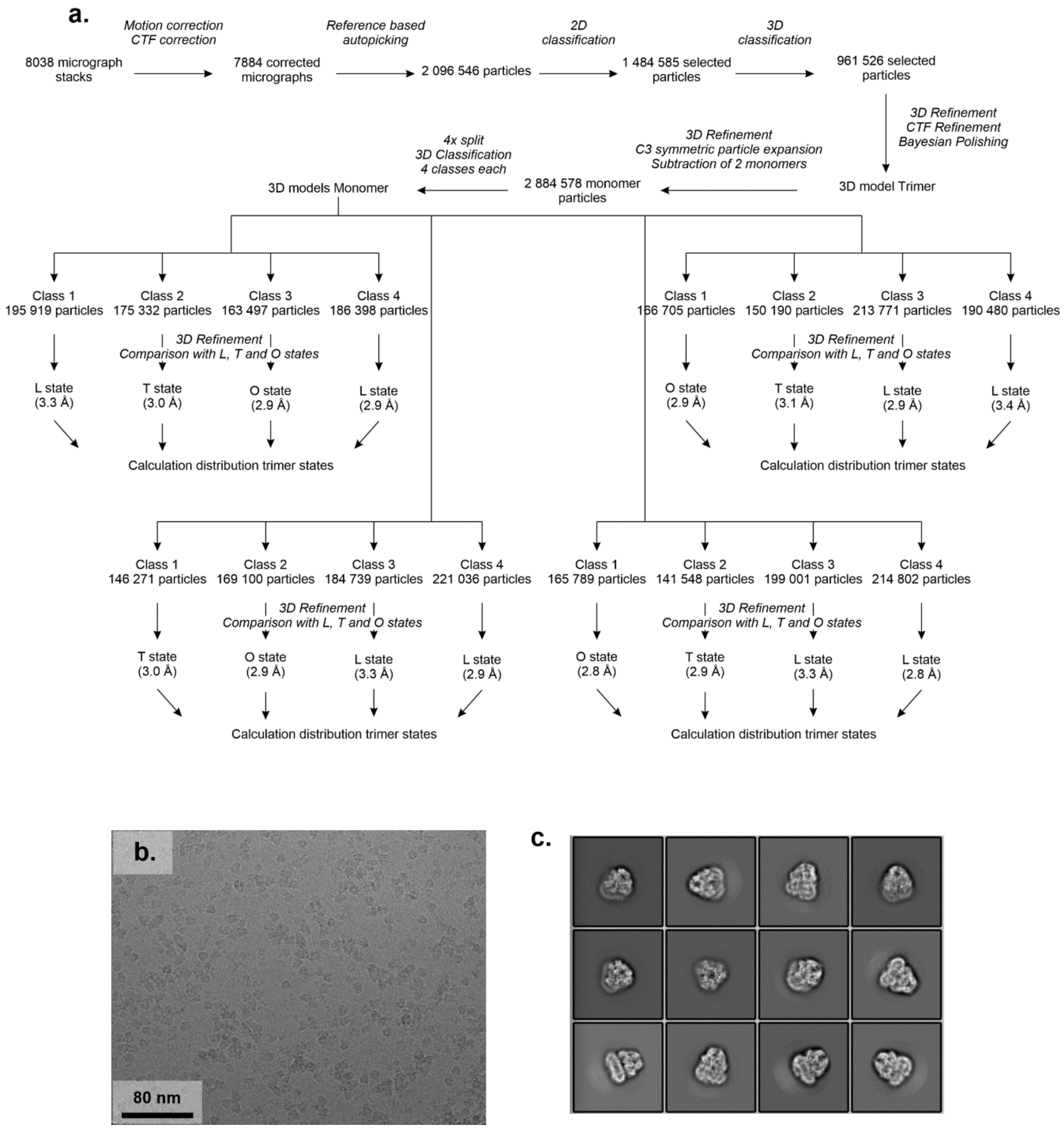


**Figure S10: Cryo-EM data of AcrB wildtype reconstituted in salipro nanodiscs. (a.)** Processing pipeline for evaluation of the distribution of conformational states. The processing was performed in Relion [3]. Due to the high number of particles the dataset was split in four parts, that were processed independently in parallel. The conformational state of each class was determined by comparison with the three monomers of the best resolved asymmetric LTO structure of AcrB wildtype (PDB ID: 4dx5). Representative micrograph and 2D classes from the dataset are shown in (**b.**). and (**c.**), respectively. The local resolution of the 3D volumes of the monomer class averages is depicted in figure S11.

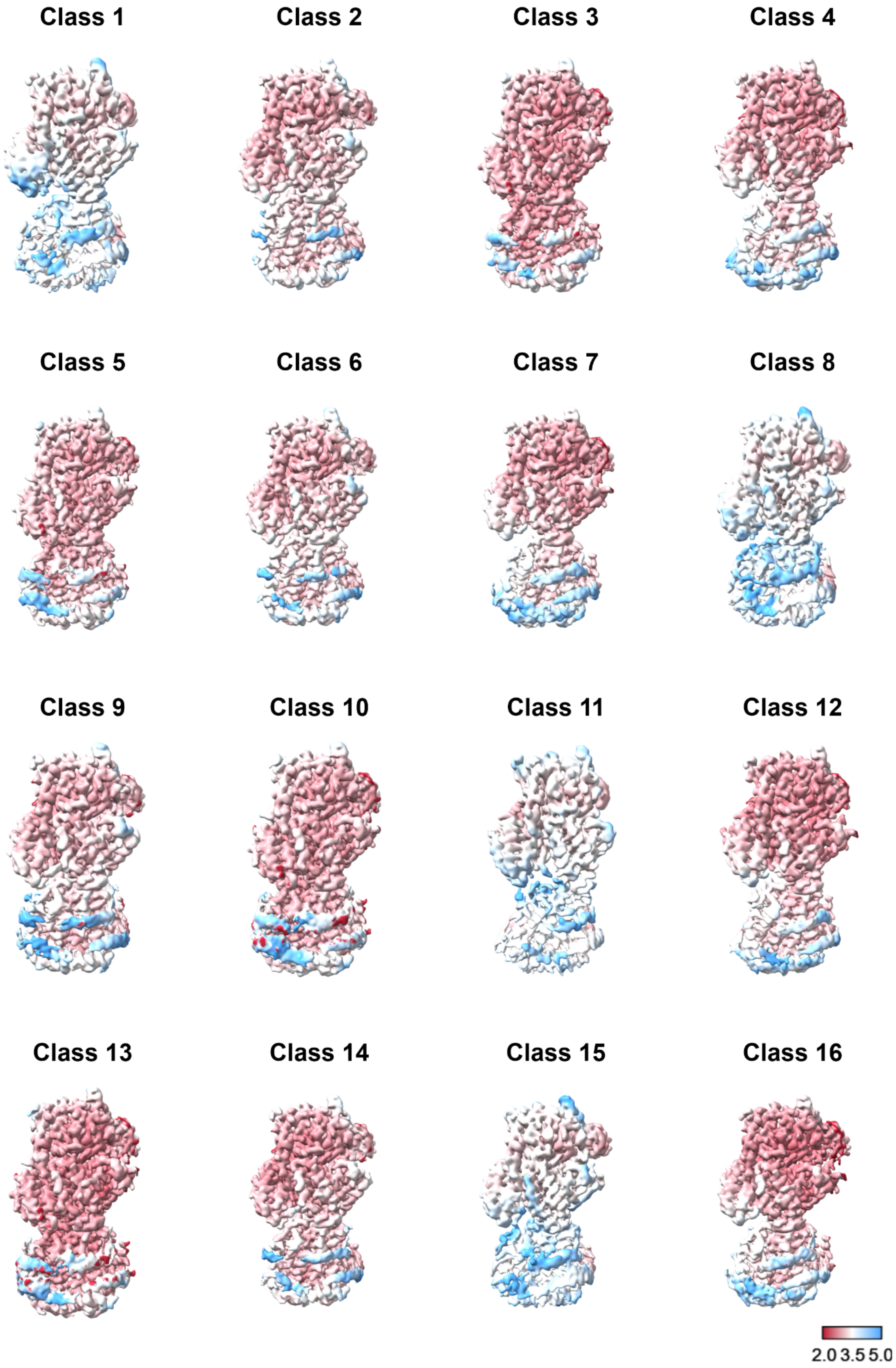


**Figure S11: Local resolution of the monomer classes from the AcrB wildtype salipro nanodiscs sample.** The monomer volumes were determined as described in figure S10.

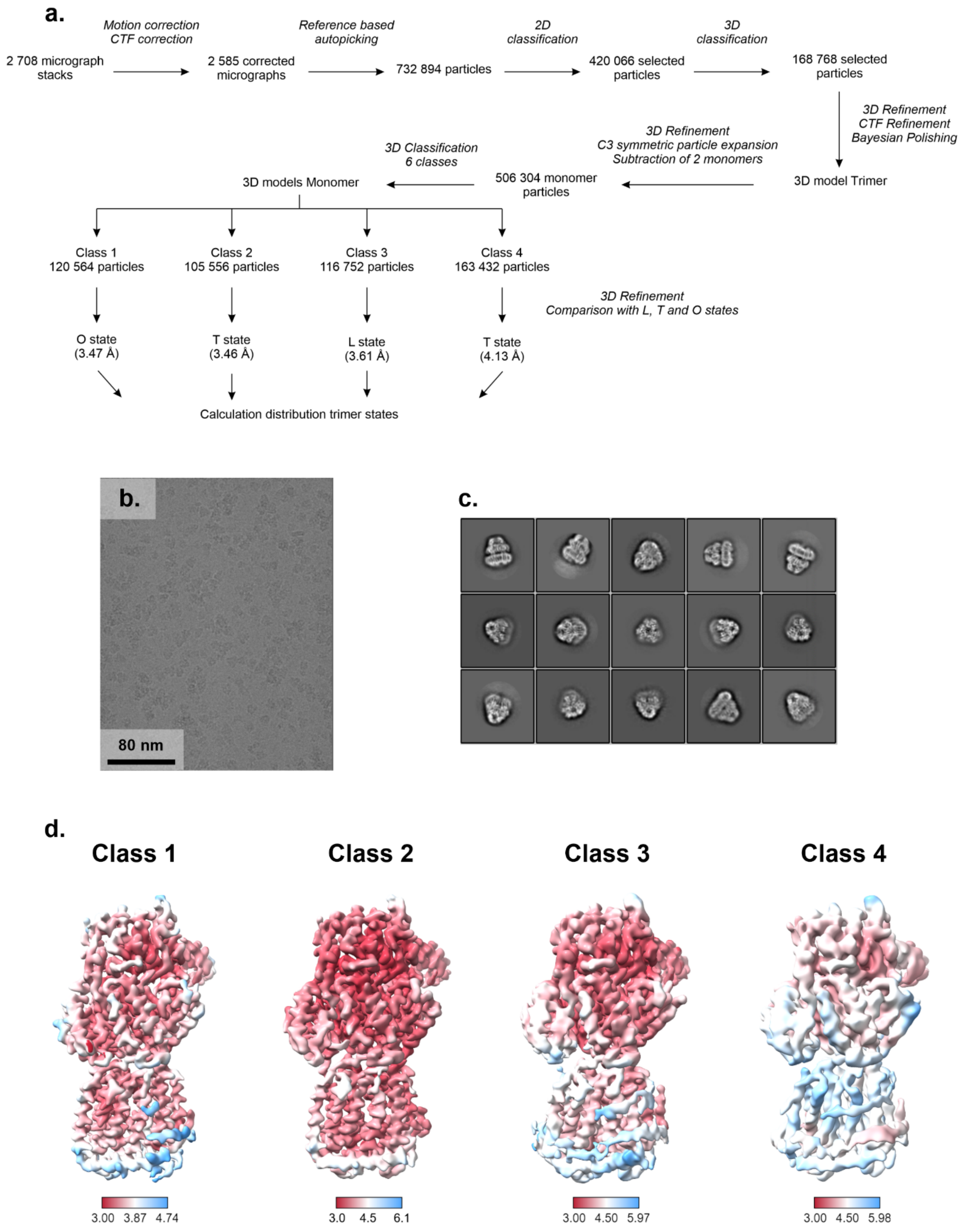


**Figure S12: Cryo-EM data of AcrB V612F reconstituted in salipro nanodiscs. (a.)** Processing pipeline for evaluation of the distribution of conformational states. The processing was performed in Relion [3]. The conformational state of each class was determined by comparison with the three monomers of the best resolved asymmetric LTO structure of AcrB wildtype (PDB ID: 4dx5). Representative micrograph and 2D classes from the dataset are shown in (**b.**). and (**c.**), respectively. The local resolution of the 3D volumes of the monomer class averages is depicted in (**d.**)

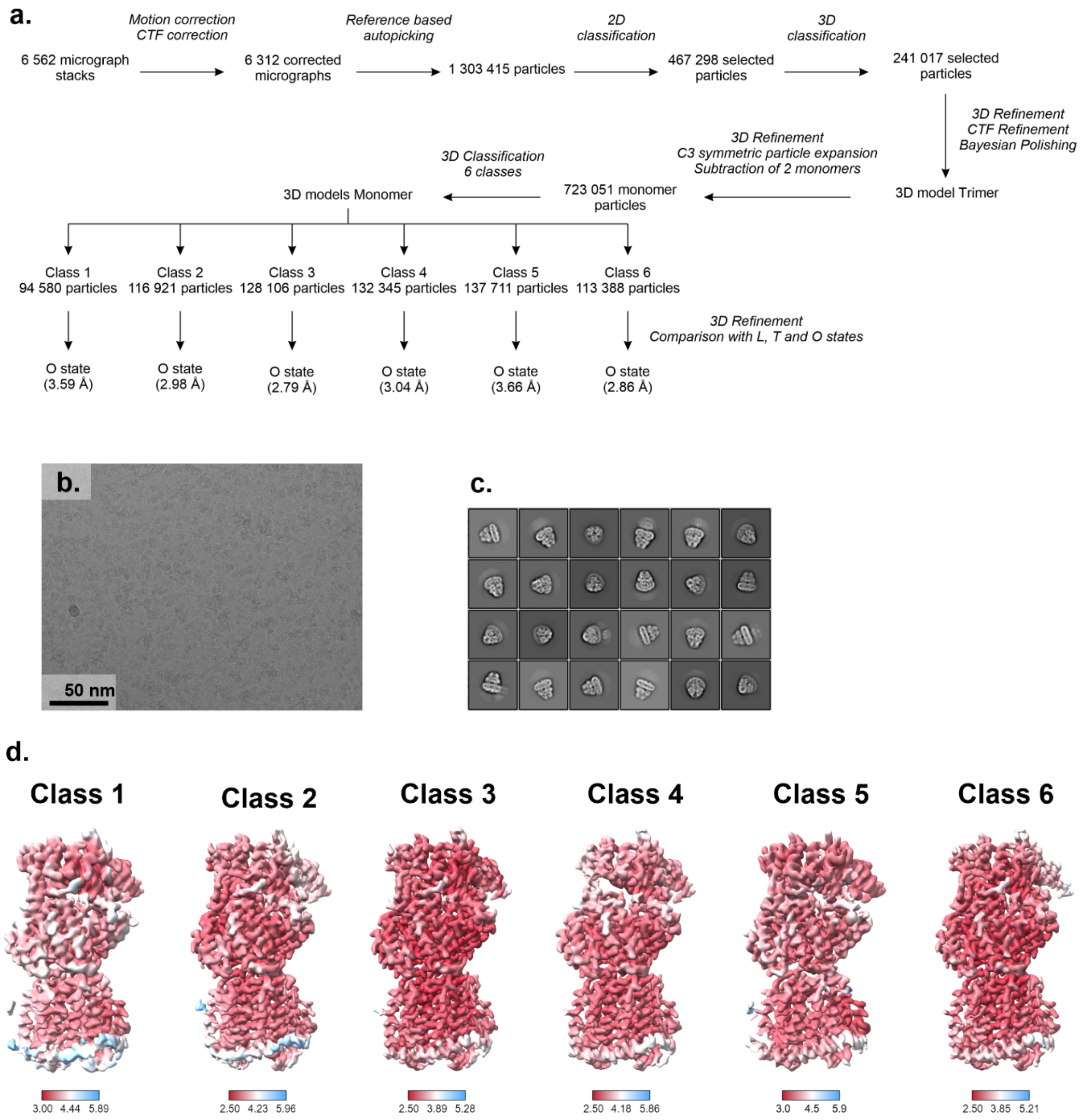


**Figure S13: Cryo-EM data of OqxB wildtype reconstituted in salipro nanodiscs. (a.)** Processing pipeline for evaluation of the distribution of conformational states. The processing was performed in Relion [3]. The conformational state of each class was determined by comparison with the three monomers of the best resolved asymmetric LTO structure of AcrB wildtype (PDB ID: 4dx5). Representative micrograph and 2D classes from the dataset are shown in (**b.**). and (**c.**), respectively. The local resolution of the 3D volumes of the monomer class averages is depicted in (**d.**)

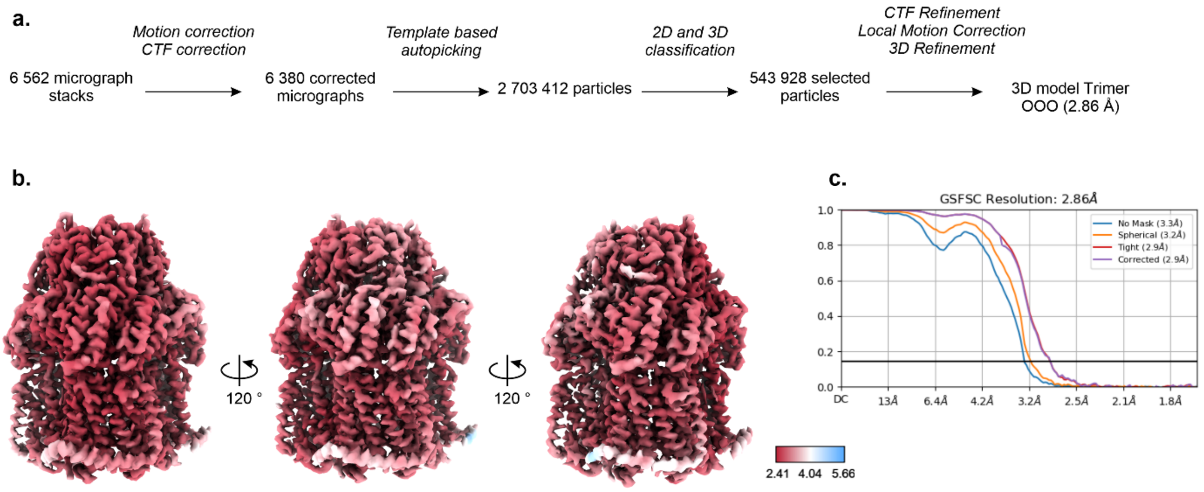


**Figure S14: Processing pipeline for structure determination of the OqxB trimer.** The same dataset as in Fig. S13 was processed again in Cryosparc [4] (**a**). The template-based picking resulted in more good particles compared to the processing in Relion [3] leading to a better resolution in the final 3D trimer map, therefore this model was used for structure building. The model was processed without imposed symmetry. The local resolution of the trimer and gold standard FSC curves are depicted in (**b.**) and (**c.**) respectively.

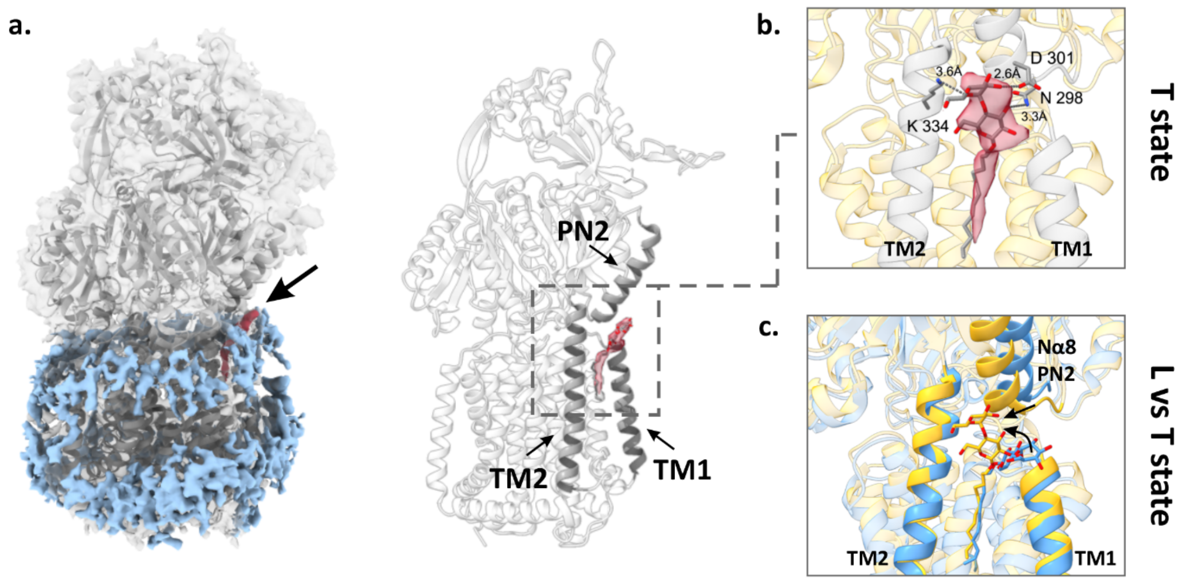


**Figure S15: DDM binding in the transmembrane domain of AcrB.** In several of the monomer classes in the cryo-EM datasets of detergent solubilised AcrB a DDM density was observed in the groove between transmembrane helices TM1 and TM2. This density is shown exemplary for class 1 of the V612W dataset. (**a**) The 3D electron density map of V612W, class 1 (left panel) was overlayed with the structure model of the AcrB T monomer with bound DDM (PDB ID: 6zoe). The micelle densities surrounding the TMD are shown in blue. The DDM molecule in the TM1/TM2 groove is depicted in red and indicated by the arrow. TM1, TM2 and the Nα8 helix from the PN2 subdomain, that coordinate the DDM molecule (red), are highlighted in the right panel. A closer zoom of the detergent binding site is depicted in (**b**). The structure model of AcrB in the T state (PDB ID: 6zoe) fitted in the density map is shown in yellow with the TM1, TM2 and Nα8 helices highlighted in grey. DDM and the residues N298, D301 and K334 that coordinate the maltose rings are depicted as sticks coloured by element with carbon atoms in grey, oxygen in red and nitrogen in blue. The DDM density from the cryo-EM map is displayed in red. (**c**) Comparison of the DDM orientation in the TM1/TM2 groove of the L and T states. The figure shows an overlay of the L (blue) and T (yellow) states of DDM bound crystallographic AcrB structures (PDB ID: 4dx5 and 6zoe). The TMD helices TM1 and TM2 and the Nα8 helix of the PN2 subdomain are indicated. DDM is shown as sticks coloured by atom type with carbon atoms in blue (L state) or yellow (T state) and oxygen atoms in red. The shifts of the maltose ring of DDM and the PN2 subdomain from the L to the T state are indicated by the arrows.

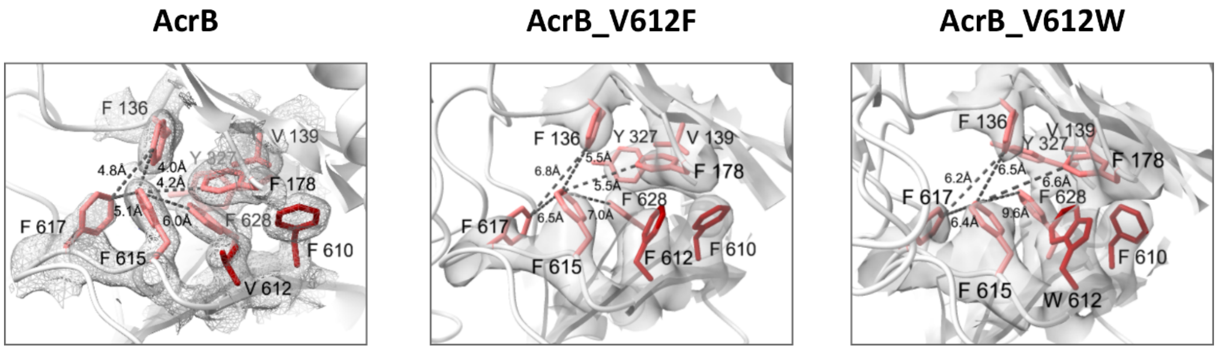


**Figure S16: Deep binding pocket (DBP) of AcrB wildtype (wt), V612F and V612W in the O state.** The conserved hydrophobic pocket residues in the DBP of AcrB wildtype (PDB ID: 4dx5) and in the structures of the best resolved V612F and V612W monomers from our cryo-EM analysis (Fig S6-S12) are shown as sticks. The residues at positions 610 and 612 are highlighted in darker red. The dashed lines indicate the distances measured between selected DBP residues. Crystallographic 2Fo-Fc density (AcrB) depicted as a mesh at σ 1. Cryo-EM maps are depicted as solid surface with contour level 0.0828 (V612F) and 0.0096 (V612W).

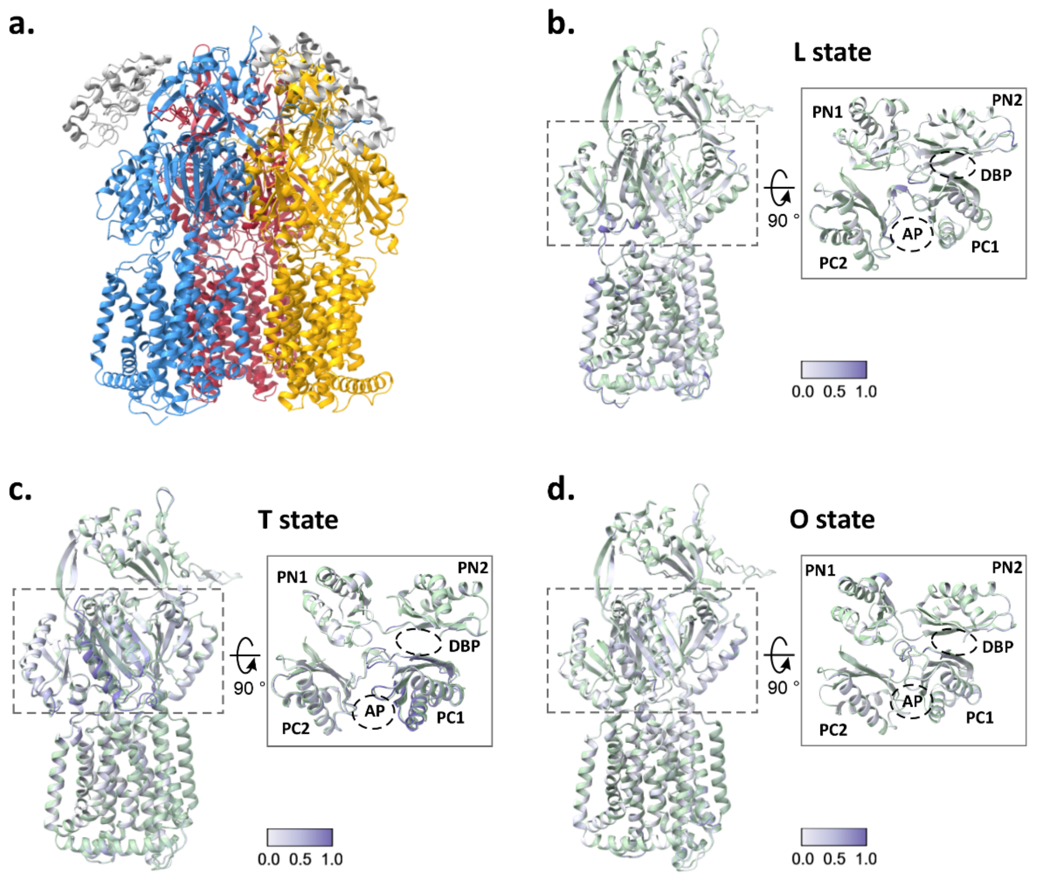


**Figure S17: Structure of AcrB V612N in the LTO state.** (**a.**) Structure of the AcrB V612N variant solved using electron density maps derived from X-ray data of crystals grown in the $P2_12_12_1$ space group. The structure was solved as an asymmetric AcrB trimer with two bound DARPins. Comparison with the asymmetric structure of AcrB wildtype (PDB ID: 4dx5) allowed to identify the V612N trimeric state as LTO. Colours: blue – L state, yellow – T state, red – O state, grey – DARPin. (**b.-d.**) Superimposition of each of the V612N monomers with the best fitting AcrB wildtype L, T, or O states is shown as side views. The V612N monomer is coloured by RMSD to the wildtype as indicated in the respective colour key, the wildtype monomer (derived from PDB ID: 4dx5) is coloured in green. Insets: top view on the AcrB porter domain with indication of the PN1, PN2, PC1, and PC2 subdomains. Dashed ovals indicate the location of the access pocket (AP) and deep binding pocket (DBP).

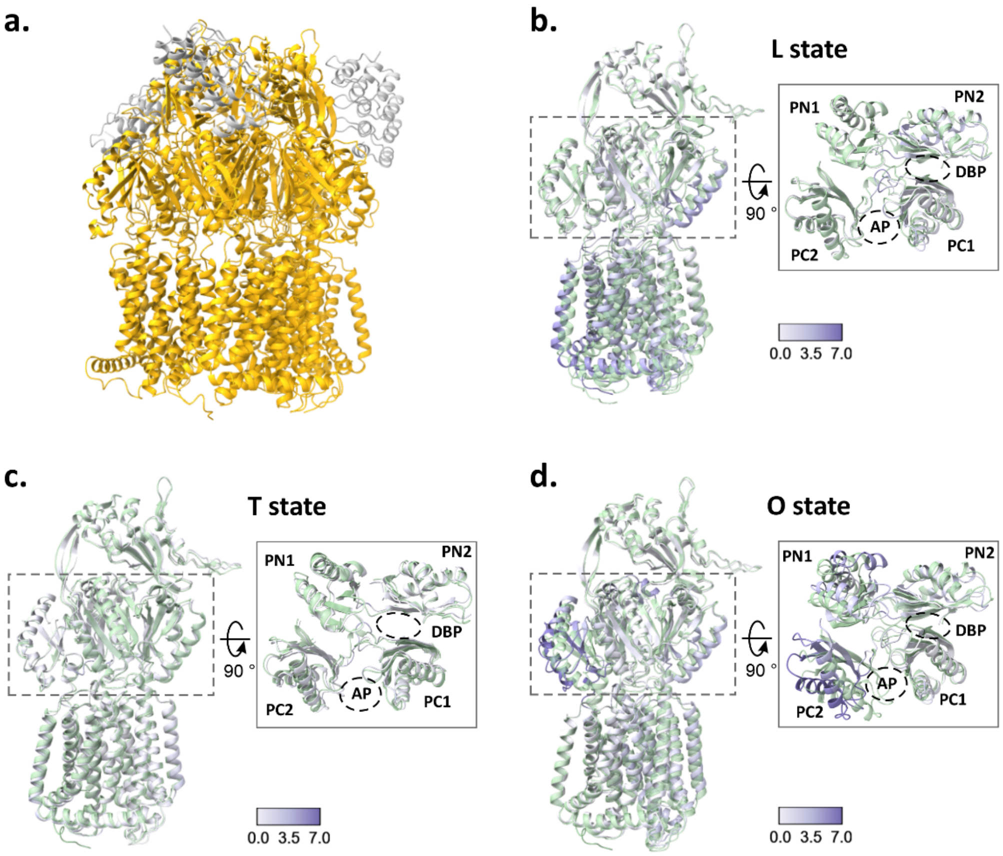


**Figure S18: Structure of AcrB V612N in the TTT state.** (**a.**) Structure of the AcrB V612N variant solved using electron density maps derived from X-ray data of crystals grown in the I23 space group. The structure was solved as a symmetric AcrB trimer with three identical chains. One DARPin molecule is associated with each chain. Comparison with the asymmetric structure of AcrB wildtype (PDB ID: 4dx5) allowed to identify the V612N trimeric state as TTT. Colours: yellow – AcrB, grey – DARPin. (**b.-d.**) Superimposition of one representative V612N monomer with each of the monomer states of the AcrB wildtype (L, T, or O) is shown as side views. The V612N monomer is coloured by RMSD to the wildtype as indicated in the respective colour key, the wildtype monomer (derived from PDB ID: 4dx5) is coloured in green. Insets: top view on the AcrB porter domain with indication of the PN1, PN2, PC1, and PC2 subdomains. Dashed ovals indicate the location of the access pocket (AP) and deep binding pocket (DBP).

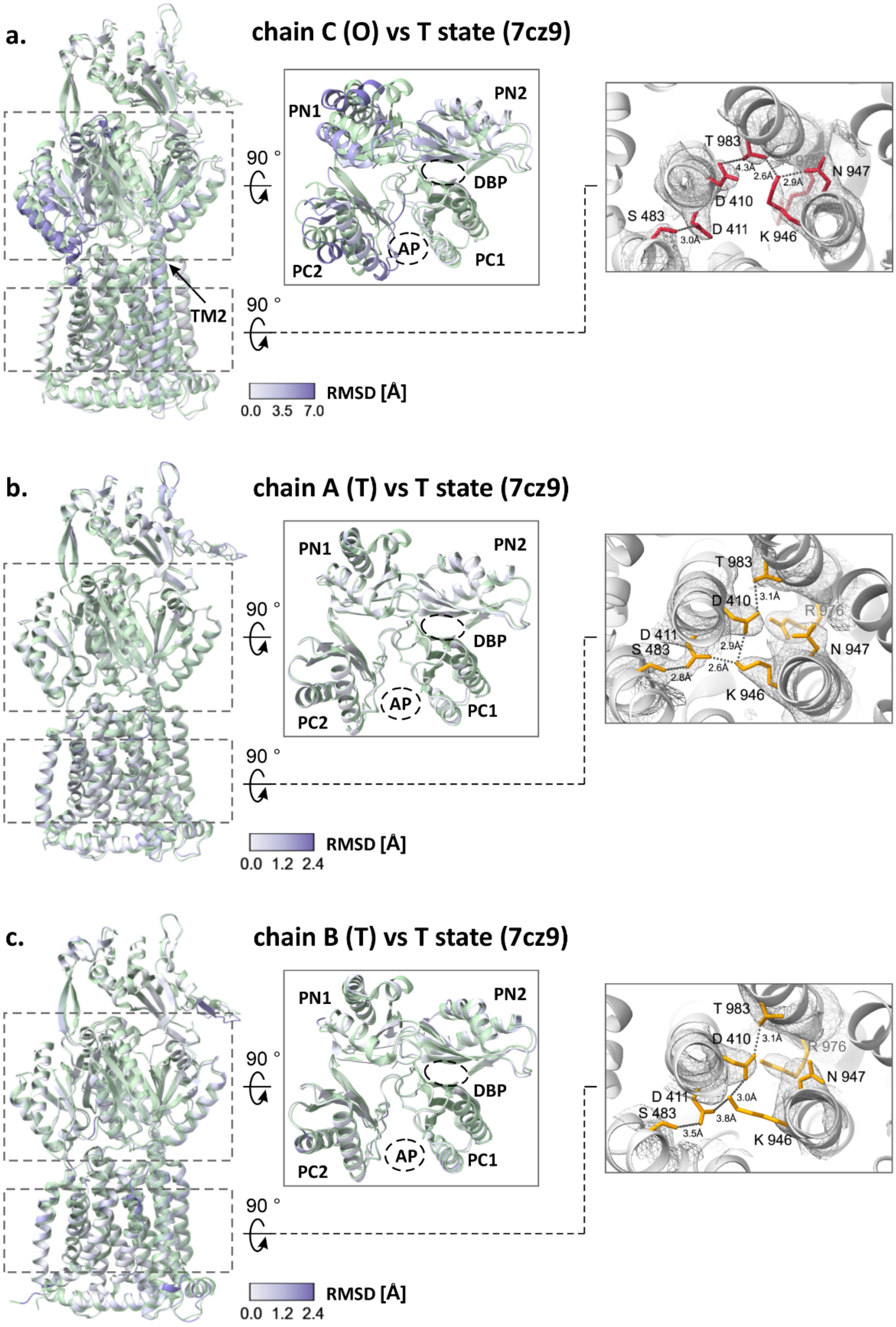


**Figure S19: Crystallographic structure of OqxB in the TTO state.** Each chain (**a-c**) of the crystallographic structure of OqxB presented in this study was overlayed with the T state of OqxB from the previously published TTT structure (PDB ID: 7cz9). Left panel: Overlay of the full-length monomers. Middle panel: a top view of the PD. The PD subdomains and the TM2 are indicated. The TTO structure is coloured by the RMSD to the T state according to the respective colour key. OqxB in the T state (PDB ID: 7cz9) is coloured green. Right panel: Proton translocation network in the TTO structure. The central titratable residues of the proton translocation network are shown as sticks in red (O state) and yellow (T states) for the TTO structure. Crystallographic 2Fo-Fc densities are shown as a mesh at σ 1. The distances between the residues are indicated by the dashed lines.

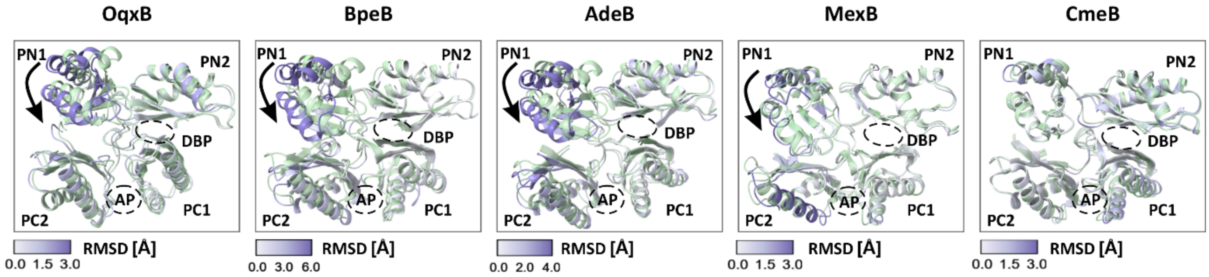


**Figure S20: Comparison of the O and O* states across RND multidrug efflux pumps.** The monomers in the O and the O* states of OqxB (this study), BpeB (PDB ID: 7wls), AdeB (PDB ID: 7kgh), MexB (PDB ID: 6ta6 and 6t7s) and CmeB (PDB ID: 5lq3) were overlayed. The O* state is coloured green, and the O state is coloured by the RMSD to O* as indicated in the respective colour key. The figure shows a top view of the porter domain. The PC1, PC2, PN1 and PN2 subdomains and the substrate binding pockets (AP, DBP) are labelled. The shift of the PN1 subdomain in the O* state is indicated by the arrows.

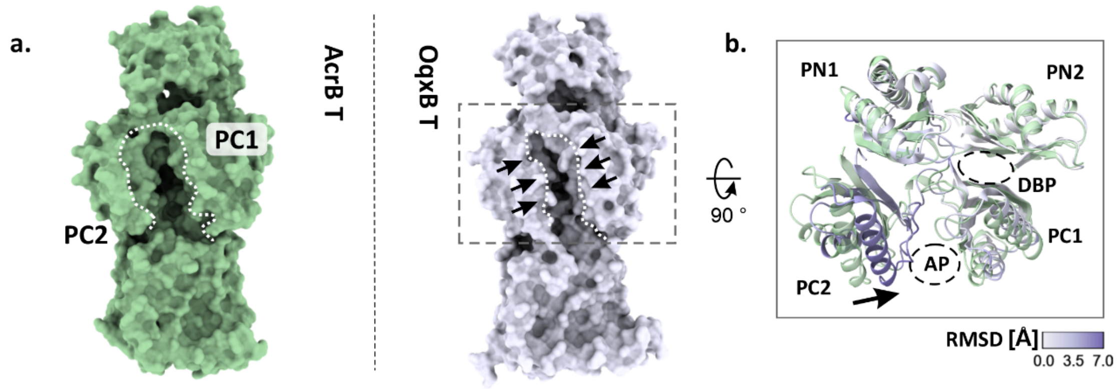


**Figure S21: Comparison of the T state of OqxB and AcrB.** (**a**) Surface representation of AcrB (PDB ID: 4dx5, left panel, green) and OqxB (PDB ID 7cz9, right panel, grey) in the T state. The entrance of the AP, flanked by the helices of the PC1 and PC2 subdomains, is indicated. Arrows indicate the reduced cleft in OqxB. The T states of AcrB and OqxB were aligned, and a top view of the porter domains is shown in (**b**). The subdomains (PN1, PN2, PC1, PC2) and the substrate binding pockets (AP, DBP) are indicated. AcrB is coloured green and OqxB is coloured by the RMSD to AcrB according to the displayed colour key. The shift of PC2 observed in OqxB is indicated by the arrow.

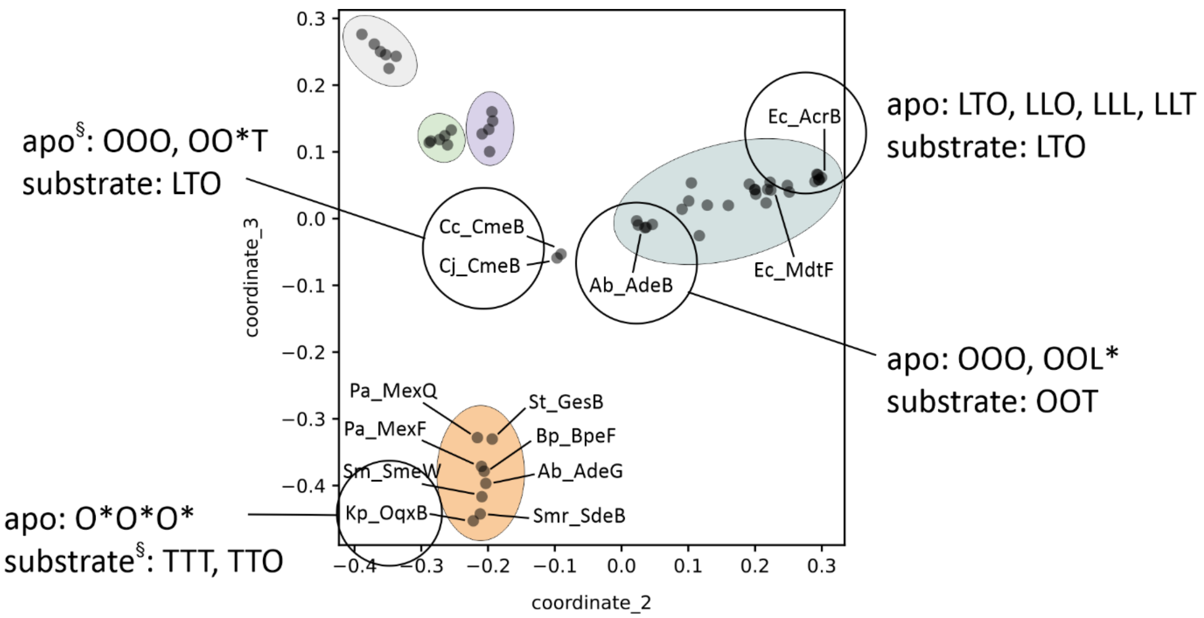


**Figure S22: Distribution of conformational states across HAE-1 RND efflux pumps.** The conformational states observed for apo and substrate bound *E. coli* AcrB (this study), *A. baumannii* AdeB [5,6], *Campylobacter* CmeB [7,8] and *K. pneumoniae* OqxB ([9] and this study) are summarised in the PaSi map from Fig. 1. The cryo-EM analysis of AcrB suggests an intrinsically flexible trimer that exists in an equilibrium between the LTO, LLL, LLT and LLO states. In contrast, AdeB that marks the far end of the AcrB cluster shows a reduced heterogeneity with only the OOO and OOL* conformations in the apo state. The L* state resembles the T state, but in contrast to T the DBP is closed. Similarly to AdeB, for apo CmeB, OOO and OO*T conformations have been described. Finally, for OqxB, a homogenous conformation with a single closed state (O*O*O*) was found for the apo protein. Differences also seem to be present in the conformations adopted in the presence of substrates. For AcrB in presence of DDM, the LTO state was predominant in the sample, whereas all further possible trimer compositions were found at low frequencies (less than 10 %). AdeB in the presence of ethidium adopted the OOT state. The LTO and OO*T states were also found in around 5% of the particles. CmeB adopted the LTO state in the presence of several different drugs. OqxB adopts the TTT and TTO states in the presence of DDM and binding of the detergent in the DBP was observed. § indicates crystallographic structures. Structural information for all other states is derived from cryo-EM studies. For the cryo-EM data, the conformations adopted by the major fraction of the particles (> 10 %) is displayed.
