## Supplementary Tables for "Conformational plasticity across phylogenetic clusters of RND multidrug efflux pumps and its impact on substrate specificity"

**Table S1: Uniprot accession numbers of the sequences used for the sequence similarity analysis.** The representative proteins of the HAE-1 RND transporter family in the transporter classification database (Saier et al. 2006) (accessed 18.08.2023) with addition of the BpeF and CmeB sequences were analysed. The protein sequences deposited in Uniprot (https://www.uniprot.org/, accessed: 18.08.2023) were used. Abbreviations: *Ec: Escherichia coli; Rr: Rhizobium radiobacter; Ab: Acinetobacter baumannii; Ng: Neisseria gonorrhoeae; Pa: Pseudomonas aeruginosa; Pp: Pseudomonas putida; Sm: Stenotrophomonas maltophilia; Ka: Klebsiella aerogenes; Bg: Burkholderia glumae; Cj: Campylobacter jejuni; Cc: Campylobacter coli; Bp: Burkholderia pseudomallei; St: Salmonella typhimurium; Vp: Vibrio parahaemolyticus; Ft: Francisella tularensis; Vc: Vibrio cholerae; Smr: Serratia marcescens; Rs: Ralstonia solanacearum; Abr: Alcanivorax borkumensis; Kp: Klebsiella pneumoniae; Ea: Erwinia amylovora; Hi: Haemophilus influenzae; Pf: Pseudomonas fluorescens*

| **Protein** | **Uniprot ID** |
|---|---|
| Ec_AcrB | P31224 |
| Ec_AcrF | P24181 |
| Rr_IfeB | O68441 |
| Ab_AdeE | Q8GKU1 |
| Ng_MtrD | Q51073 |
| Pa_MexB | P52002 |
| Ec_AcrD | P24177 |
| Pp_ArpB | Q9KJC2 |
| Pp_TtgB | O52248 |
| Pp_TtgE | Q9KWV4 |
| Pp_TtgH | Q93PU4 |
| Ec_MdtB | P76398 |
| Ec_MdtC | P76399 |
| Ec_MdtF | P37637 |
| Sm_SmeW | B2FLY4 |
| Pa_MexD | Q51396 |
| Pa_MexF | Q9I0Y8 |
| Pa_MexK | Q9HXW4 |
| Pf_EmhB | Q6V6X8 |
| Ka_EefB | Q8GC83 |
| Bg_ToxH | Q4VSJ4 |
| Pa_MexY | Q9ZNG8 |
| Cj_CmeB | Q8RTE4 |
| Cc_CmeB | A0A1L2IWC1 |

| | |
|---|---|
| Bp_BpeB | Q6VV68 |
| Bp_AmrB | O87936 |
| St_GesB | Q8ZRG9 |
| Vp_VmeB | Q2AAU3 |
| Pa_TriC | Q9I6X4 |
| Ft_AcrB | A0Q8A5 |
| Ab_AdeJ | Q24LT7 |
| Vc_VexF | A6P7H3 |
| Smr_SdeB | Q84GI9 |
| Pa_MexI | Q9HWH4 |
| Pa_MexW | A0A6A9K223 |
| Pa_MexQ | Q4LDT6 |
| Pa_MexN | Q4LDT8 |
| Vc_VexB | Q9KVI2 |
| Vc_VexD | A6P7H1 |
| Vc_VexK | Q9KRG9 |
| Pa_MuxC | Q9I0V7 |
| Pa_MuxB | Q9I0V6 |
| Ab_AdeB | Q2FD70 |
| Sm_SmeB | Q9RBY8 |
| Sm_SmeE | Q9F240 |
| Sm_SmeJ | A0A0U5D5D4 |
| Ab_AdeG | Q2FD81 |
| Rs_AcrB | Q8Y3H0 |
| Abr_AcrB | Q0VQY6 |
| Kp_OqxB | C5IZH1 |
| Ea_AcrB | E3DBE3 |
| Kp_AcrB | Q93K40 |
| Hi_AcrB | Q57124 |
| Bp_BpeF | Q63NK6 |

**Table S2: AcrB substrates for the phenotype assays.** The table lists the AcrB substrates used in the plate dilution assays (PDA) and for minimal inhibitory concentration (MIC) determination. The substrate class, the abbreviation used in the figures and the concentration used in the PDAs are indicated.

| **Substrate** | **Class** | **Abbreviation** | **Concentration [µg/mL]** |
|---|---|---|---|
| Chloramphenicol | Phenicol | CAM | 1.5 |
| Thiamphenicol | Phenicol | TIA | 12 |
| Linezolid | Oxazolidinone | LIN | 20 |
| Tetracycline | Tetracycline | TET | 0.3 |
| Oxytetracycline | Tetracycline | OxTET | 0.25 |
| Tigecycline | Tetracycline | TIG | 0.125 |
| Chlortetracycline | Tetracycline | ClTET | 0.2 |
| Doxycycline | Tetracycline | DXC | 0.5 |
| Minocycline | Tetracycline | MIN | 0.3 |
| Dicloxacillin | Beta-Lactam (Penicillin) | DIC | 36 |
| Oxacillin | Beta-Lactam (Penicillin) | OXA | 16 |
| Piperacillin | Beta-Lactam (Penicillin) | PIP | 0.075 |
| Dodecyl-β-D-maltosid | Detergent | DDM | 75 |
| Sodium dodecyl sulfate | Detergent | SDS | 70 |
| Erythromycin | Macrolide | ERY | 20 |
| Clarythromycin | Macrolide | CLA | - |
| Novobiocin | Aminocoumarin | NOV | 10 |
| Fusidic acid | Steroid | FUA | 12 |
| Doxorubicin | Anthracycline | DOX | 40 |
| Hoechst33342 | Dye | H33342 | 5 |
| Tetraphenylphosphonium | Phosphonium cation | TPP | 300 |

**Table S3: Minimal inhibitory concentration (MIC) values.** BW25133 Δ*acrB* cells expressing AcrB wildtype (wt) or V612 variants were exposed to a serial dilution of a toxic substrate. The inactive D407N was used as negative control. The MIC value corresponds to the first dilution step at which no growth was detected. The table shows the mean MIC value for at least three biological replicates in µg/mL.

| | **wt** | **D407N** | **V612F** | **V612W** | **V612N** | **V612A** |
|---|---|---|---|---|---|---|
| **Chloramphenicol** | 4 | 1-2 | 8 | 8 | 8 | 8 |
| **Thiamphenicol** | 32 | 16 | 64 | 32-64 | 64 | 32 |
| **Linezolid** | 128 | 8 | 256 | 256 | 128-256 | 256 |
| **Tetracycline** | 1 | 0.25-0.5 | 1 | 1 | 1 | 1 |
| **Oxytetracycline** | 0.5 | 0.25 | 1 | 1 | 1 | 1 |
| **Tigecycline** | 0.25 | 0.125 | 0.125-0.25 | 0.25 | 0.25 | 0.25 |
| **Chlortetracycline** | 1 | 0.25 | 1 | 1 | 1 | 1 |
| **Doxycycline** | 2 | 0.25 | 2 | 2 | 2 | 2 |
| **Minocycline** | 4-16 | 0.25-8 | 0.5-2 | 16 | 16 | 16 |
| **Dicloxacillin** | 256 | 8-32 | 128 | 128 | 128 | 256 |
| **Oxacillin** | 128 | 2-8 | 64 | 64-128 | 64 | 128 |
| **Piperacillin** | 0.5 | 0.0625-0.125 | 0.125-0.25 | 0.25-0.5 | 0.25-0.5 | 0.5 |
| **SDS** | >8192 | 64 | 64 | 64-128 | >8192 | >8192 |
| **Erythromycin** | 128 | 8 | 64 | 64 | 64 | 64-128 |
| **Clarythromycin** | 64-128 | 4-16 | 32-64 | 64 | 32 | 64 |
| **Novobiocin** | 8-16 | 1-4 | 2-8 | 8 | 8 | 16-32 |
| **Fusidic acid** | 64-128 | 8-16 | 32-64 | 32-64 | 32 | 64 |
| **Hoechst33342** | 4-8 | 1-2 | 2-4 | 2 | 4 | 4 |

**Table S4: Estimated binding free energies for the docking poses in figure S5.** ΔGb gives the total binding free energy **in kcal/mol.** The contributions from each individual residue involved in the interactions are indicated. Values are coloured from highest (white) to lowest (grey). Abbreviations: MIN – minocycline, DOX- doxorubicin, ERY – erythromycin, CAM – chloramphenicol.

**MIN**

| | *ΔGb* | *S48* | *Q176* | *L177* | *F178* | *G179* | *S180* | *E273* | *N274* | *I277* | *A279* | *S287* | *F610* | *V/F/W612* | *F615* | *R620* |
|---|---|---|---|---|---|---|---|---|---|---|---|---|---|---|---|---|
| **wt** | -28.2 | -0.7 | | -1.4 | -6.3 | -4.2 | -1.3 | -1.5 | -5.7 | -5.8 | -0.7 | -0.6 | -0.8 | -1.5 | -2.4 | -7.8 |
| **V612F** | -28.7 | -1.0 | -1.1 | -3.0 | -6.4 | -7.1 | -2.6 | | -4.3 | -5.2 | -1.0 | | -0.6 | -2.1 | -1.0 | |
| **V612W** | -25.6 | -3.5 | -0.7 | -1.8 | -5.8 | -4.0 | -2.3 | -1.3 | -5.3 | -5.0 | -0.7 | | | -2.3 | -1.5 | |

**DOX**

| | *ΔGb* | *T44* | *S46* | *Q89* | *S128* | *E130* | *S132* | *S133* | *S134* | *F136* | *Q176* | *F178* | *G179* | *I277* | *F610* | *V/F/W612* | *F615* | *F617* | *R620* | *F628* |
|---|---|---|---|---|---|---|---|---|---|---|---|---|---|---|---|---|---|---|---|---|
| **wt** | -53.5 | -1.1 | -5.9 | -3.8 | -6.7 | -7.8 | | | | -1.3 | -5.2 | -5.5 | | -1.9 | -1.0 | -2.0 | -2.3 | -0.7 | | |
| **V612F** | -46.4 | | -2.2 | -3.4 | -2.2 | -4.9 | | | -1.1 | | -5.8 | -7.8 | -1.2 | -3.4 | -0.8 | -4.0 | -3.1 | | -1.8 | -1.1 |
| **V612W** | -54.2 | -2.9 | | -5.1 | | -5.3 | -7.7 | -1.7 | -5.7 | -2.1 | -3.6 | -1.9 | | | | -3.5 | -5.9 | -1.0 | | |

**ERY**

| | *ΔGb* | *T44* | *Q89* | *T91* | *E130* | *S132* | *S133* | *S134* | *F136* | *Q176* | *F178* | *K292* | *V/F/W612* | *F615* | *F617* | *R620* | *E673* |
|---|---|---|---|---|---|---|---|---|---|---|---|---|---|---|---|---|---|
| **wt** | -50.3 | -2.0 | -7.0 | -2.4 | -2.5 | -7.7 | -2.4 | -12.1 | -2.2 | -3.8 | -1.7 | -5.7 | | -5.2 | -3.6 | -1.8 | -2.1 |
| **V612F** | -52.1 | -2.1 | -6.9 | -2.3 | -2.5 | -7.6 | -2.6 | -12.1 | -2.1 | -3.8 | -1.7 | -5.6 | -2.2 | -5.2 | -3.6 | -1.7 | -2.1 |
| **V612W** | -52.8 | -2.1 | -6.9 | -2.3 | -2.5 | -7.6 | -2.6 | -12.1 | -2.1 | -3.8 | -1.6 | -5.6 | -2.9 | -5.5 | -3.6 | -1.6 | -2.1 |

**CAM**

| | *ΔGb* | *S134* | *F136* | *V139* | *Q151* | *Q176* | *L177* | *F178* | *G179* | *I277* | *I278* | *A279* | *P326* | *Y327* | *V571* | *M573* | *F610* | *V/F/W612* | *F615* | *F617* | *I626* | *F628* | *L668* | *V672* |
|---|---|---|---|---|---|---|---|---|---|---|---|---|---|---|---|---|---|---|---|---|---|---|---|---|
| **wt** | -29.6 | -7.1 | -3.0 | | | | | | | | | | | -3.3 | -1.7 | -2.4 | | | -1.5 | -1.1 | -0.6 | -5.4 | -1.5 | -1.9 |
| **V612F** | -33.6 | | | | -1.4 | -4.6 | -3.7 | -6.1 | -2.8 | -4.9 | -0.8 | -1.6 | | | | | -0.7 | -3.6 | -1.8 | | -0.8 | -1.0 | | |
| **V612W** | -30.6 | | | -1.4 | | | | -2.3 | | -1.4 | | | -0.9 | -1.5 | | -0.8 | | -8.1 | -5.3 | -1.4 | -1.4 | -4.7 | | |

**Table S5: Structural comparison of the TTO structure of OqxB (8zxs, this study) with the TTT structure of OqxB (7cz9) and LTO structure of AcrB (4dx5)**

| | | AcrB(4dx5) | | | OqxB (7cz9) | | |
|---|---|---|---|---|---|---|---|
| | | Chain A (L) | Chain B (T) | Chain C (O) | Chain A (T) | Chain B (T) | Chain C (T) |
| OqxB (8zxs) | Chain A | 2.95 | 2.81 | 3.50 | 0.81 | 1.06 | 0.89 |
| | Chain B | 3.27 | 2.69 | 3.49 | 0.85 | 0.92 | 0.99 |
| | Chain C | 3.72 | 3.77 | 2.39 | 2.51 | 2.40 | 2.49 |

The RMSD (root mean square deviation) in Å is calculated between aligned pairs of the backbone Cα atoms in each monomer by LSQKAB in CCP4 program suite.

**Table S6: Data collection and refinement statistics crystallographic structures AcrB V612W**

| | AcrB V612W MIY (TTT) PDB 9FE2 | AcrB V612W apo (TTT) PDB 9FE3 |
|---|---|---|
| **Data collection** | | |
| Space group | I 2 3 | I 2 3 |
| Cell dimensions | | |
| *a*, *b*, *c* (Å) | 227.49, 227.49, 227.49 | 227.43, 227.43, 227.43 |
| α, β, γ (°) | 90.00, 90.00, 90.00 | 90.00, 90.00, 90.00 |
| Resolution (Å) | 46.44 - 1.89 (1.96 - 1.89) | 44.60 - 2.30 (2.38 - 2.30) |
| $R_{sym}$ or $R_{merge}$ | 0.01656 (1.971) | 0.01593 (2.029) |
| $I / \sigma I$ | 21.91 (0.32) | 19.98 (0.33) |
| Completeness (%) | 96.99 (70.06) | 97.68 (77.33) |
| Redundancy | 2.0 (2.0) | 2.0 (2.0) |
| **Refinement** | | |
| Resolution (Å) | 46.44 - 1.89 (1.96 - 1.89) | 44.60 - 2.3 (2.38 - 2.3) |
| No. reflections | 150429 (10838) | 84315 (6626) |
| $R_{work}$ / $R_{free}$ | 0.2239 / 0.2455 | 0.2535 / 0.2909 |
| No. atoms | | |
| Protein | 9499 | 9073 |
| Ligand/ion | 33 | 0 |
| Water | 384 | 6 |
| *B*-factors | | |
| Protein | 56.84 | 97.59 |
| Ligand/ion | 83.56 | - |
| Water | 53.54 | 65.66 |
| R.m.s. deviations | | |
| Bond lengths (Å) | 0.012 | 0.003 |
| Bond angles (°) | 1.19 | 0.65 |

*Values in parentheses are for highest-resolution shell.

**Table S7: Data collection and refinement statistics crystallographic structures AcrB V612F**

| | AcrB V612F MIY (TTT)<br>PDB 9FHC | AcrB V612F apo (TTT)<br>PDB 9FE4 |
|---|---|---|
| **Data collection** | | |
| Space group | I 2 3 | P 3 2 1 |
| Cell dimensions | | |
| *a*, *b*, *c* (Å) | 227.46, 227.46, 227.46 | 134.41, 134.41, 190.97 |
| α, β, γ (°) | 90.00, 90.00, 90.00 | 90.00, 90.00, 120.00 |
| Resolution (Å) | 29.87 - 2.20 (2.279 - 2.20) | 49.70 - 2.80 (2.90 - 2.80) |
| $R_{sym}$ or $R_{merge}$ | 0.1293 (0.9247) | 0.09319 (3.095) |
| $I / \sigma I$ | 27.16 (4.31) | 20.19 (0.90) |
| Completeness (%) | 94.51 (67.04) | 99.73 (99.79) |
| Redundancy | 24.9 (21.4) | 11.0 (10.3) |
| **Refinement** | | |
| Resolution (Å) | 29.87 - 2.20 (2.279 - 2.20) | 49.70 - 2.80 (2.90 - 2.80) |
| No. reflections | 93142 (6569) | 49611 (4859) |
| $R_{work}$ / $R_{free}$ | 0.2080 / 0.2380 | 0.2696 / 0.3030 |
| No. atoms | | |
| Protein | 9115 | 7843 |
| Ligand/ion | 33 | 0 |
| Water | 281 | 0 |
| *B*-factors | | |
| Protein | 40.76 | 110.83 |
| Ligand/ion | 39.25 | - |
| Water | 33.70 | - |
| R.m.s. deviations | | |
| Bond lengths (Å) | 0.008 | 0.009 |
| Bond angles (°) | 1.27 | 1.21 |

*Values in parentheses are for highest-resolution shell.

**Table S8: Data collection and refinement statistics crystallographic structures AcrB V612N**

| | AcrB V612N (LTO)<br>PDB 9FHG | AcrB V612N (TTT)<br>PDB 9FHJ |
|---|---|---|
| **Data collection** | | |
| Space group | P 21 21 21 | I 2 3 |
| Cell dimensions | | |
| $a$, $b$, $c$ (Å) | 145.81, 161.40, 245.41 | 228.65, 228.65, 228.65 |
| $\alpha$, $\beta$, $\gamma$ (°) | 90.00, 90.00, 90.00 | 90.00, 90.00, 90.00 |
| Resolution (Å) | 49.44 - 3.00 (3.11 - 3.00) | 41.74 - 3.55 (3.68 - 3.55) |
| $R_{sym}$ or $R_{merge}$ | 0.06477 (0.8325) | 0.01899 (2.092) |
| $I / \sigma I$ | 6.86 (1.03) | 15.67 (0.33) |
| Completeness (%) | 99.75 (99.71) | 93.89 (45.09) |
| Redundancy | 2.0 (2.0) | 2.0 (2.0) |
| **Refinement** | | |
| Resolution (Å) | 49.44 - 3.00 (3.11 - 3.00) | 41.74 - 3.55 (3.68 - 3.55) |
| No. reflections | 115982 (11502) | 22677 (1078) |
| $R_{work}$ / $R_{free}$ | 0.2239 / 0.2829 | 0.2719 / 0.3260 |
| No. atoms | | |
| Protein | 25972 | 9086 |
| Ligand/ion | 0 | 0 |
| Water | 0 | 0 |
| *B*-factors | | |
| Protein | 79.72 | 195.91 |
| Ligand/ion | - | - |
| Water | - | - |
| R.m.s. deviations | | |
| Bond lengths (Å) | 0.009 | 0.004 |
| Bond angles (°) | 1.14 | 0.68 |

*Values in parentheses are for highest-resolution shell.

**Table S9: Data collection and refinement statistics crystallographic structure OqxB**

| | OqxB (TTO)<br>PDB 8ZXS |
|---|---|
| **Data collection** | |
| Space group | P 21 21 21 |
| Cell dimensions | |
| *a*, *b*, *c* (Å) | 121.39, 165.94, 249.03 |
| α, β, γ (°) | 90.00, 90.00, 90.00 |
| Resolution (Å) | 49.34 - 2.75 (2.80 - 2.75) |
| $R_{sym}$ or $R_{merge}$ | 0.082 (>1.0) |
| $I / \sigma I$ | 12.9 (0.9) |
| Completeness (%) | 99.65 (95.19) |
| Redundancy | 7.33 (7.30) |
| **Refinement** | |
| Resolution (Å) | 49.34 - 2.75 |
| No. reflections | 130594 (6157) |
| $R_{work}$ / $R_{free}$ | 0.2363 / 0.2877 |
| No. atoms | |
| Protein | |
| Ligand/ion | |
| Water | |
| *B*-factors | 124,7 |
| Protein | |
| Ligand/ion | |
| Water | |
| R.m.s. deviations | |
| Bond lengths (Å) | 0,005 |
| Bond angles (°) | 0,888 |

*Values in parentheses are for highest-resolution shell.

**Table S10: Cryo-EM data collection, refinement and validation statistics**

| | OqxB (O*O*O*)<br>EMD-50334<br>PDB 9FDZ | AcrB V612F (O)<br>EMD-50332<br>PDB 9FDQ | AcrB V612W (O)<br>EMD-50331<br>PDB 9FDP |
|---|---|---|---|
| **Data collection and processing** | | | |
| Magnification | 105000 | 130000 | 105000 |
| Voltage (kV) | 300 | 300 | 300 |
| Electron exposure (e–/Å$^2$) | 50 | 60 | 50 |
| Defocus range (μm) | -0.8 to -2.4 | -0.5 to -3.0 | -0.8 to -3.5 |
| Pixel size (Å) | 0.837 | 0.68 | 0.837 |
| Symmetry imposed | no | no | no |
| Initial particle images (no.) | 2703412 | 732894 | 856118 |
| Final particle images (no.) | 543928 | 120564 | 81754 |
| Map resolution (Å) | 2.86 | 3.47 | 3.3 |
| FSC threshold | 0.143 | 0.143 | 0.143 |
| **Refinement** | | | |
| Initial model used (PDB code) | AlphaFold | 4dx5 | 4dx5 |
| Model resolution (Å) | 2.86 | 3.4 | 3.3 |
| FSC threshold | 0.143 | 0.143 | 0.143 |
| Map sharpening *B* factor (Å$^2$) | - | - | - |
| Model composition | | | |
| Non-hydrogen atoms | 23736 | 7853 | 7857 |
| Protein residues | 3120 | 1033 | 1033 |
| Ligands | 0 | 0 | 0 |
| *B* factors (Å$^2$) | | | |
| Protein | 49.18/217.07/114.41 | 14.15/112.77/38.75 | 14.15/112.77/38.75 |
| Ligand | - | - | - |
| R.m.s. deviations | | | |
| Bond lengths (Å) | 0.004 | 0.002 | 0.002 |
| Bond angles (°) | 0.951 | 0.549 | 0.507 |
| Validation | | | |
| MolProbity score | 1.10 | 1.18 | 1.15 |
| Clashscore | 3.10 | 3.91 | 3.59 |
| Poor rotamers (%) | 0.59 | 0.00 | 0.00 |
| Ramachandran plot | | | |
| Favored (%) | 98.78 | 99.22 | 99.32 |
| Allowed (%) | 1.22 | 0.78 | 0.68 |
| Disallowed (%) | 0.00 | 0.00 | 0.00 |
