## Supplementary Information for "Conformational plasticity across phylogenetic clusters of RND multidrug efflux pumps and its impact on substrate specificity"

**Homology modelling, ensemble-docking, and free binding energy calculations**

Despite our efforts to determine experimental co-structures with different AcrB substrates bound to the V612F/W variants, we only obtained co-structures with minocycline. To assess changes in the DBP interactions for further substrates, we therefore performed a computational study, namely docking and free energy calculations. For our study we choose four established AcrB substrates: minocycline, doxorubicin, erythromycin and chloramphenicol. For minocycline experimental co-structures are present for the wildtype (PDB ID: 4dx5) and the V612F/W variants (this study). This allows us to evaluate how the experimental data correlate with the computational study. For doxorubicin an experimental co-structure is present for the wildtype (PDB ID: 4dx7) which also gives us a cross-reference for the computed results. Doxorubicin is an anthracycline and its chemical structure and binding site within AcrB (deep binding pocket, DBP) is very similar to the tetracycline antibiotics. However, in contrast to the tetracyclines, doxorubicin shows much higher reduction in the resistance activity of the variants. Further, there is a distinct phenotype discrepancy between the V612F and V612W variants (Fig. 1 main manuscript). For erythromycin and chloramphenicol, the V612F/W substitution induced changes of the phenotype that mimic the phenotype of the MdtF variant and of the proteins from the OqxB cluster, which contain a F at the position equivalent to V612 [1–6].

The top docking binding poses of minocycline closely resemble the experimental structures (Supplementary, Fig. S5). Moreover, the results of the free binding energy calculation suggest that minocycline is stabilised by a network of interactions involving the hydrophilic groups of the ligand and the polar side chains or backbone carbonyl and amide groups of S48, L177, G179, S180, N274, I277, and A279 (Supplementary, table S4). F178, F610, F615 and the F/W612 also contribute to the interaction. In wildtype AcrB, R620 greatly contributes to the free binding energy, but this interaction is lost in the V612F and V612W variants as observed also in the experimental structures. However, this is compensated by a higher contribution of the substituted F/W612, as well as further individual side chains (L177, G179, S180 for V612F and S48, S148 for V612W). The total binding free energy for the variants is similar to the wildtype with a difference of only 0.5 kcal/mol for V612F and 2.6 kcal/mol for V612W. Thus, the calculated binding poses and the experimental data both suggest the same mode of minocycline binding with hydrophilic interactions between the polar groups of the ligand and polar residues and the backbone of AcrB, and coordination of the aromatic ring of minocycline between F178, F615 and F/W612. In agreement with the experimental data, the docking calculations confirm that the loss or decrease of the individual interactions due to the shift in the binding position of minocycline in V612F/W are readily compensated by alternative interactions with further residues in the binding site.

The top binding pose calculated for doxorubicin in the DBP of wildtype AcrB closely resembles the orientation observed in the previously published experimental structure of the wildtype [7] (Supplementary, Fig. S5) despite a slight (~1.5 Å) sliding back towards the entrance of the DBP. A similar but much more pronounced shift (> 4.5 Å) is observed for doxorubicin in V612W. In V612F, the calculated pose of the ligand is slightly tilted compared to the experimental structure. The differences in the orientation of V612F compared to the wildtype also induces changes in the coordination network, and many of the interactions observed for the wildtype are weakened or abolished in V612F (e.g. with T44, S46, S128, E130, F136) (Supplementary, table S4). Even though this is compensated by stronger or additional interactions with S134, F178, G179, I277, F612, F615, R620 and F628, the total free binding energy for V612F is 7.1 kcal/mol higher than the wildtype. For V612W, the interactions are weaker compared to the wildtype for the same amino acid sidechains as for V612F. In contrast to V612F, for V612W this is compensated mainly by polar interactions with the serine-rich loop (S132, S133, S134), and with T44 and Q89, owned to the shift in the position of the ligand in the V612W structure. The difference in the total binding energy for V612W compared to the wildtype (0.7 kcal/mol) is much lower than for V612F.

Similarly to minocycline, in the docking poses for doxorubicin a sliding of the ligand is observed for V612W and indeed the overlay with the experimental doxorubicin structure shows a clear steric overlap of the ligand with the W612 side chain (Supplementary, Fig. S5). However, in the V612F docking results, such shift in the ligand position is not observed. Instead, here the F612 side chain is flipped away from the ligand binding site. As it can be seen from the overlay of the experimental V612F structure with the doxorubicin docking pose, F612 in the experimental structure would clash with the ligand (Fig. SI1). The difference in the orientations of F612 in the experimental and docking structures might represent alternative conformations that F612, and potentially W612, can adopt.

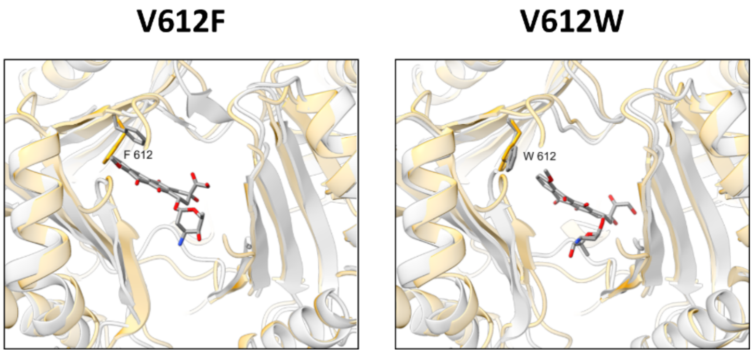


**Figure SI1: Overlay of the experimental structures of AcrB V612F and V612W with the doxorubicin docking poses.** The figure shows the top docking pose of doxorubicin in the DBP of the V612F and V612W variants. AcrB is coloured grey with F/W612 and doxorubicin shown as sticks and coloured by atom type with carbon – grey, oxygen – red, and nitrogen – blue. The docking results are overlayed with the experimental crystallographic structures of AcrB V612F and V612W (this study) coloured in yellow.

The experimental and computational results for the binding of minocycline and the computational results for the binding of doxorubicin suggest that the steric hindrance introduced by the V612F/W substitution results in a sliding of the substrate towards the entrance of the DBP. Besides minocycline, one further representative of the tetracycline class was found to bind in the same position in the DBP [8] and it is feasible that other tetracyclines bind in a similar fashion as well. Thus, it is likely that similar sliding of the drug occurs, but the versatility of the DBP presumably allows accommodation of the substrate and formation of an alternative interaction network, explaining the marginal change in the resistance phenotype of V612F/W against minocycline and other tetracyclines (see Fig. 1 main manuscript).

Despite the discussed differences in the binding poses of minocycline and doxorubicin, their binding sites are very similar in AcrB wildtype, V612F and V612W (Supplementary, Fig. S5). In contrast, the top binding poses of chloramphenicol differ greatly in the three proteins (Fig. SI2a). Chloramphenicol is one of the smallest AcrB ligands and with a van-der-Waals volume of 249 Å$^3$ (calculated with Chemicalize, https://chemicalize.com/) it is almost 15x smaller than the volume of the DBP (approximately 3700 Å$^3$ [9]). The ligand is small, flexible and amphipathic, and it is likely that it can be coordinated in different grooves within the DBP. Molecular dynamics (MD) simulations have shown that chloramphenicol frequently flips in the DBP of the T state [10,11], suggesting that this substrate can be accommodated in different poses and frequently changes between them. A cryo-EM structure of AcrB in the presence of chloramphenicol showing a density for the ligand in the DBP has been reported [10]. An overlay of the electron density map of this structure with the docking poses for the wildtype, V612F and V612W shows that the putative ligand density is in proximity of the docking pose for the wildtype (Fig. SI2b).

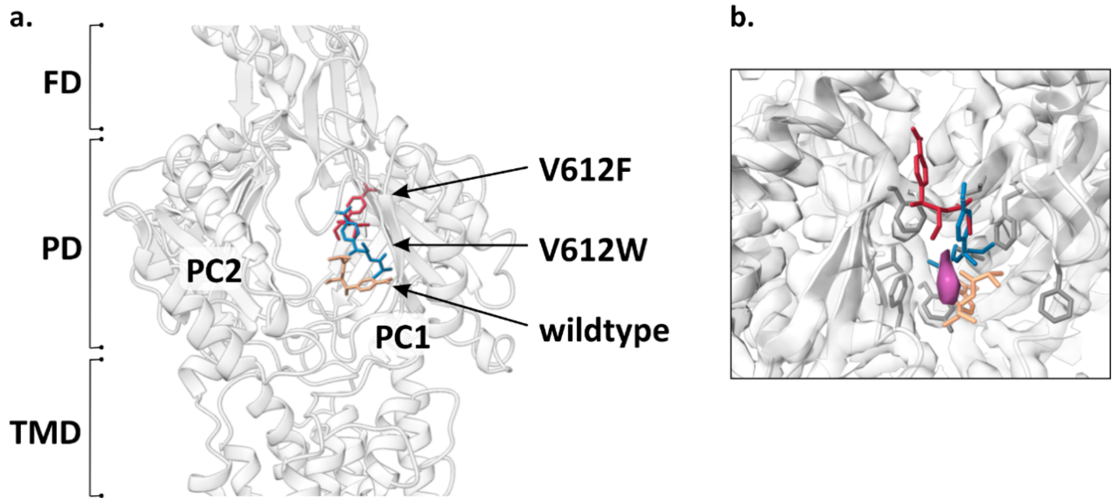


**Figure SI21: Chloramphenicol binding in the DBP of AcrB.** (**a**) The top docking poses for chloramphenicol in the DBP of AcrB wildtype, V612F and V612W. The AcrB structure is outlined in the cartoon representation. Chloramphenicol docking poses are shown as sticks and coloured yellow for the wildtype, red for V612F and blue for V612W. Abbreviations: FD – funnel domain, PD – porter domain, TMD – transmembrane domain. The PC1 and PC2 subdomains are indicated. (**b**) Overlay of the top docking poses for chloramphenicol with the experimental AcrB wildtype structure in the presence of this substrate (PDB ID: 6sgr). The docking poses for the wildtype, V612F and V612W are shown as sticks coloured as in (a). The cryo-EM electron density map of the experimental structure is shown as surface with the putative chloramphenicol density highlighted in purple. The residues of the DBP are shown as sticks in grey.

The different computational poses might indicate distinct binding sites within the DBP. The reason why one of these binding sites is preferred in the wildtype and the others in the variants, might be connected to the introduced substitution. In both V612F and V612W, the ligand is pulled up in the DBP and oriented so that the aromatic ring of chloramphenicol is facing the F/W612 side chain. Particularly in V612W it is well evident how the aromatic ring of the ligand is sandwiched between F178, F615 and W612 and these interactions likely stabilise the binding at this position. In the F612 variant the ligand similarly engages in aromatic interactions with F178 and F612 (Supplementary, Fig. S5 and table S4).

For chloramphenicol, a slight increase in the resistance conferred by V612F and V612W was observed (see Fig. 1 main manuscript). One can speculate that chloramphenicol can diffuse in and out of the AP and DBP while interacting with multiple low affinity binding sites in the PD. The additional aromatic interactions with F/W612 could stabilise the interactions with the hydrophobic cluster, pulling chloramphenicol further into the DBP and increasing its retention time, thus allowing a more efficient transport. Similar consideration might apply for further flexible, amphipathic, low molecular weight substrates that contain an aromatic ring. One AcrB substrate that fits this description is linezolid, and for this substrate a slight increase in the resistance conferred by V612F/W was observed as well.

Experimental co-structures of erythromycin show binding of the substrate at the interface between the AP and the DBP in the L state of *E. coli* AcrB [12–14]. In the close homolog AcrB from *K. pneumoniae* (96 % sequence similarity to *E. coli* AcrB), erythromycin binding within the DBP in the T state has been described [15] (Fig. SI3a). Presumably, this substrate initially binds in the L state and is guided to the interior of the porter domain during the transition from L to T. The docking pose of erythromycin in AcrB wildtype (T state) is in closer proximity to the DBP compared to the experimental co-structure of erythromycin bound to the L state (Fig. SI3a). However, the docking position of erythromycin does not reach the same binding site as the seen in the experimental structure of *K. pneumoniae* AcrB in the T state. As the calculated pose is located between both binding sites seen in the experimentally derived co-structures, it might represent an intermediate interaction mode along the way from the AP towards the DBP (Fig. SI3a).

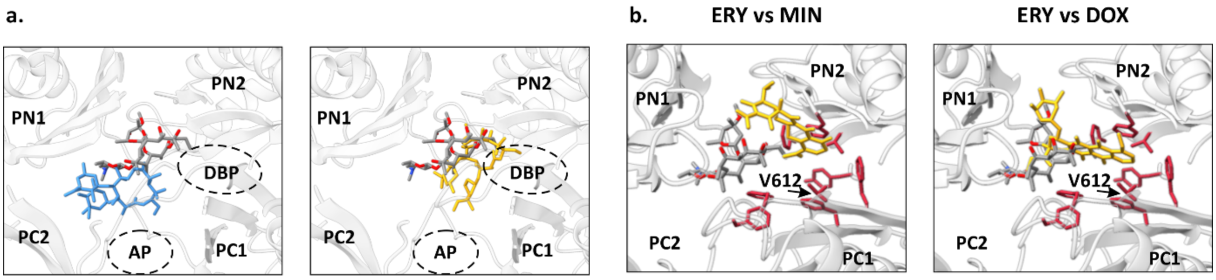


**Figure SI3: Erythromycin binding sites in the PD of AcrB.** (**a**) In experimental co-structures of *E. coli* AcrB, erythromycin binds at the interface of the AP and DBP in the L state monomer (blue, left panel, PDB ID: 3aoc). In the close homolog AcrB from *K. pneumoniae*, the substrate was found in the DBP in the T state monomer (yellow, right panel, PDB ID: 8ffs). The top docking pose for AcrB wildtype (coloured by atom type with carbon – grey, oxygen – red, and nitrogen – blue) was overlayed with both structures. The PD is outlined as cartoon and the PN1, PN2, PC1 and PC2 subdomains as well as the AP and DBP are indicated. (**b**) An overlay of the docking pose of erythromycin with the minocycline (MIN, left panel) and doxorubicin (DOX, right panel) structures. A top view of the porter domain is shown in grey and the PN1, PN2, PC1 and PC2 subdomains are indicated. The residues of the DBP are shown as sticks in red and V612 is indicated. The top docking pose of erythromycin in wildtype AcrB is shown as sticks coloured by atom type with carbon – grey, oxygen – red, and nitrogen – blue. Minocycline and doxorubicin from the experimental structures (PDB ID: 4dx5 and 4dx7 respectively) are shown as sticks in yellow.

The docking pose of erythromycin is near the entrance of the DBP, but it does not reach the DBP groove in contrast to minocycline and doxorubicin (Fig. SI3b). The latter two bind deeply within the hydrophobic groove cluster and are in immediate vicinity of the residue at position 612. Erythromycin, however, is located further away from this residue. Therefore, the steric effects of the V612F/W substitution, that were observed for minocycline and doxorubicin, are likely less relevant for erythromycin. Further, erythromycin does not contain an aromatic ring for interaction with the introduced aromatic substitution like e.g. chloramphenicol. Considering the greater distance to and minor interactions with the residue at position 612, it is possible that the V612F/W substitutions have a less pronounced effect on the interaction network of erythromycin compared to the other investigated substrates. In agreement with these considerations, the docking calculations showed identical binding poses for erythromycin in AcrB wildtype, V612F and V612W and similar binding energies with only a minor difference due to the contribution from F/W612 (Supplementary, Fig. S5 and table S4).

As the experimental structures of erythromycin show binding of this substrate in the AP-DBP-interface in the L state, it is plausible that the drug is sequestered from the periplasm through the AP in this state. Likely the substrate is then guided to the PD interior during the transition from L to T and finally reached the DBP in the T state. Potentially, the initial interactions require the PD architecture in the L state and are less favourable if the AP adopts the T state structure, as proposed earlier[11]. In contrast to the wildtype, the V612 variants exhibits a structure with increased abundance of the T state at the expense of the L state. The reduced amount of L state monomers for initial binding might compromise the transport of substrates such as erythromycin, that require the interactions for initial uptake in this state.

### Methods

Ensemble-docking calculations were performed for all compounds on the DBP of the T state ($DBP_T$) of AcrB (lined by residues S46, Q89, S128, E130, S134, F136, V139, Q176, L177, F178, G179, S180, E273, N274, D276, I277, A279, A286, P326, Y327, M573, F610, V/F/W612, F617, R620, F628, F664, F666, L668, P669, V672 [16]) using the software GNINA [17]. Following a previous approach [18,19], we generated a set of AcrB (wt, V612F, and V612W) structures featuring the largest structural variance at the $DBP_T$, in order to account for minor to medium structural changes at this site, which notoriously influence the outcome of docking calculations [20]. The list of AcrB structures we employed includes the following PDB IDs: 2dhh, 2dr6, 2drd, 2gif, 2hrt, 2j8s, 3aoa, 3aob, 3aoc, 3aod, 3noc, 3nog, 3w9h, 4dx5, 4dx6, 4dx7, 4u8v, 4u8y, 4u95, 4u96, 4zit, 4ziv, 4zjl, 5jmn, 5nc5, 5yil, 6q4n, 6q4o, 6q4p. In addition, for the V612F and the V612W AcrB variants, we included the new experimental structures reported here. Note that some of the afore listed structures contain gaps, modified residues, and mutations. Therefore, we generated consistent structures of the wt as well as the two variants of AcrB by means of homology modelling calculations, using the software MODELLER 10.2 [21]. More precisely, for the wt structures only the missing parts were modelled, using the wildtype structure (PDB ID: 4dx5) as a template. The models of the mutant structures were generated by modelling only the amino acids within 5 Å from the mutated residues, using as templates the experimental V612F and V612W structures. In both cases, the list of residues was the following: F178, I278, F610, A611, F/W612, N613, F615, G625, I626, F628. The rest of the structure was not modified, apart for gaps and additional mutations. Therefore, a multi-template homology modelling was performed for both the wt and the mutant structures. Moreover, we introduced the F/W612 mutations within the asymmetric LTO structures based on the TTT experimental geometries of V612F/W. Before running homology modelling calculations, all structures were aligned to the $DBP_T$ of the structure with PDB ID 4dx5, and only the amino acids from 1 to 1030 (in each chain) were used to ensure that all the models will have the same length.

After generating a pool of AcrB structures identical in sequence, we reassign protonation states of the protein following previous literature [9,22]. Next, we aligned all structures (29 for wt and 31 for mutants) to the $DBP_T$ of that identified by the PDB ID 4dx5 and calculated 29x29 (wt) and 31x31 (mutants) matrices of pairwise RMSDs values. From these matrices, we retained only the structures displaying RMSDs (calculated on all the heavy atoms of $DBP_T$) larger than 1 Å from each other. For pairs with RMSDs values below this threshold, we removed the structure with the lowest resolution from the pool.

The following structure data files (.sdf) were used to prepare the pdb files for docking with GNINA: CAM and ERY were downloaded from our antibiotic database (structure optimized by means of quantum-mechanical calculations) [23], while MIN and DOX were taken from the PDB database (IDs 4dx5 and 4dx7, respectively). For each ligand, the dominant protonation states at physiological pH were assigned by ChemAxon's Marvin suite of program. Docking was performed centring the grid on $DBP_T$ and using the following parameters: exhaustiveness: 512 (default 8); nmodes: 10.

Poses were first filtered according to the Convolutional Neural Networks (CNN) score provided by GNINA (which ranges from 0 – very unlikely pose – to 1 – very likely that the pose represents a true structure), retaining only those with a value larger than 2/3. Next, these poses were ranked according to their estimated affinity to the binding site.

Estimation of the binding free energy were obtained through the MM/GBSA approach [24] implemented in the MMPBSA.py tool of AMBER22 [25], following the same protocol used in previous studies [26,27]. This approach provides an intrinsically simple method for decomposing the free energy of binding into contributions from single atoms and residues, as well as ligand-residue pairs [28]. The solute conformational entropy contribution (TΔSconf) was not evaluated [24]. Calculations were performed on the top pose of each complex.
