## Supplementary material for "Conformational plasticity across phylogenetic clusters of RND multidrug efflux pumps and its impact on substrate specificity": Source Data

Biological replicate 1, technical replicate 1

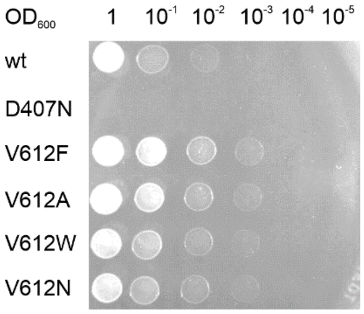


*Chloramphenicol [1.5 µg/mL]*

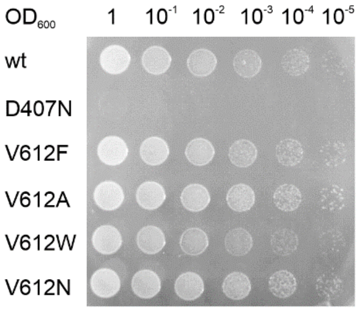


*Thiamphenicol [12 µg/mL]*

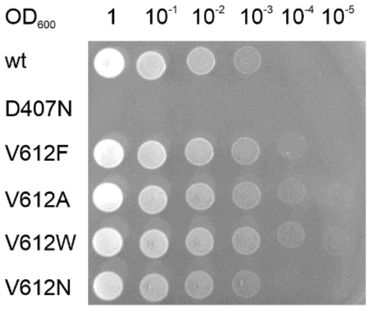


*Linezolid [20 µg/mL]*

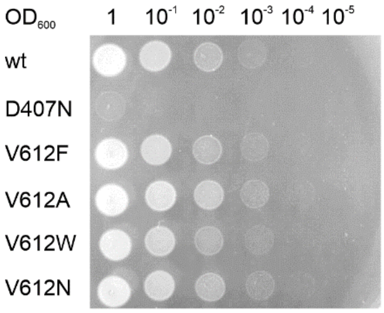


*Tetracycline [0.3 µg/mL]*

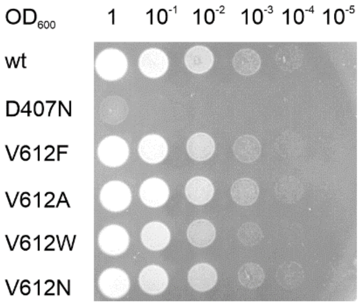


*Oxytetracycline [0.25 µg/mL]*

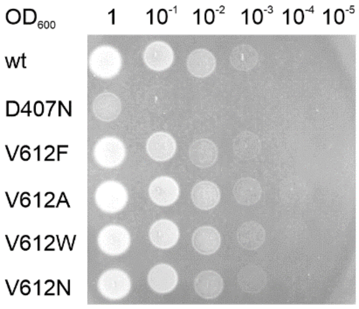


*Tigecycline [0.125 µg/mL]*

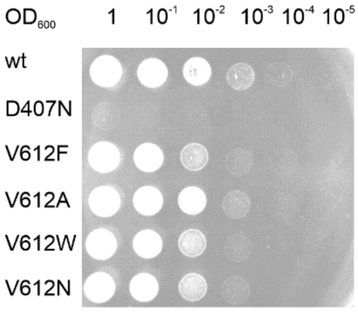


*Chlortetracycline [0.2 µg/mL]*

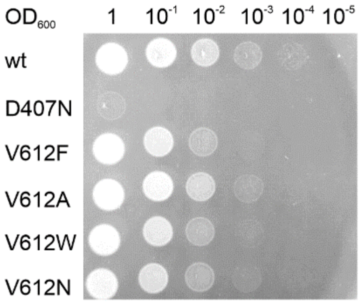


*Doxycycline [0.5 µg/mL]*

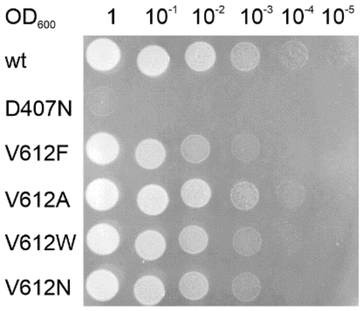


*Minocycline [0.3 µg/mL]*

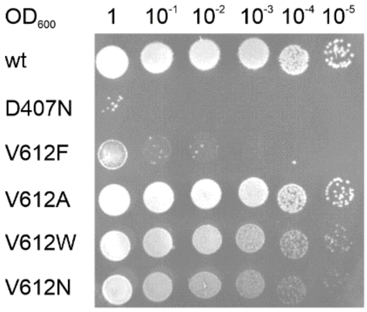


*Dicloxacillin [36 µg/mL]*

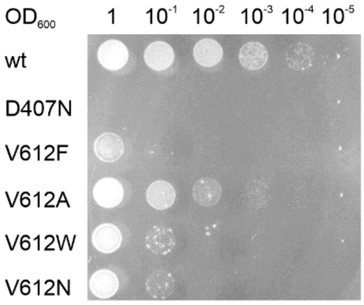


*Oxacillin [16 µg/mL]*

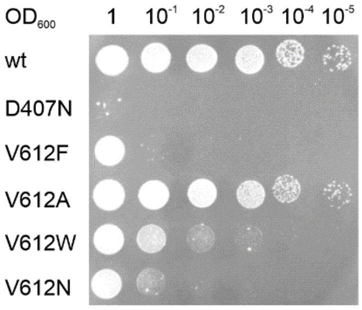


*Piperacillin [0.075 µg/mL]*

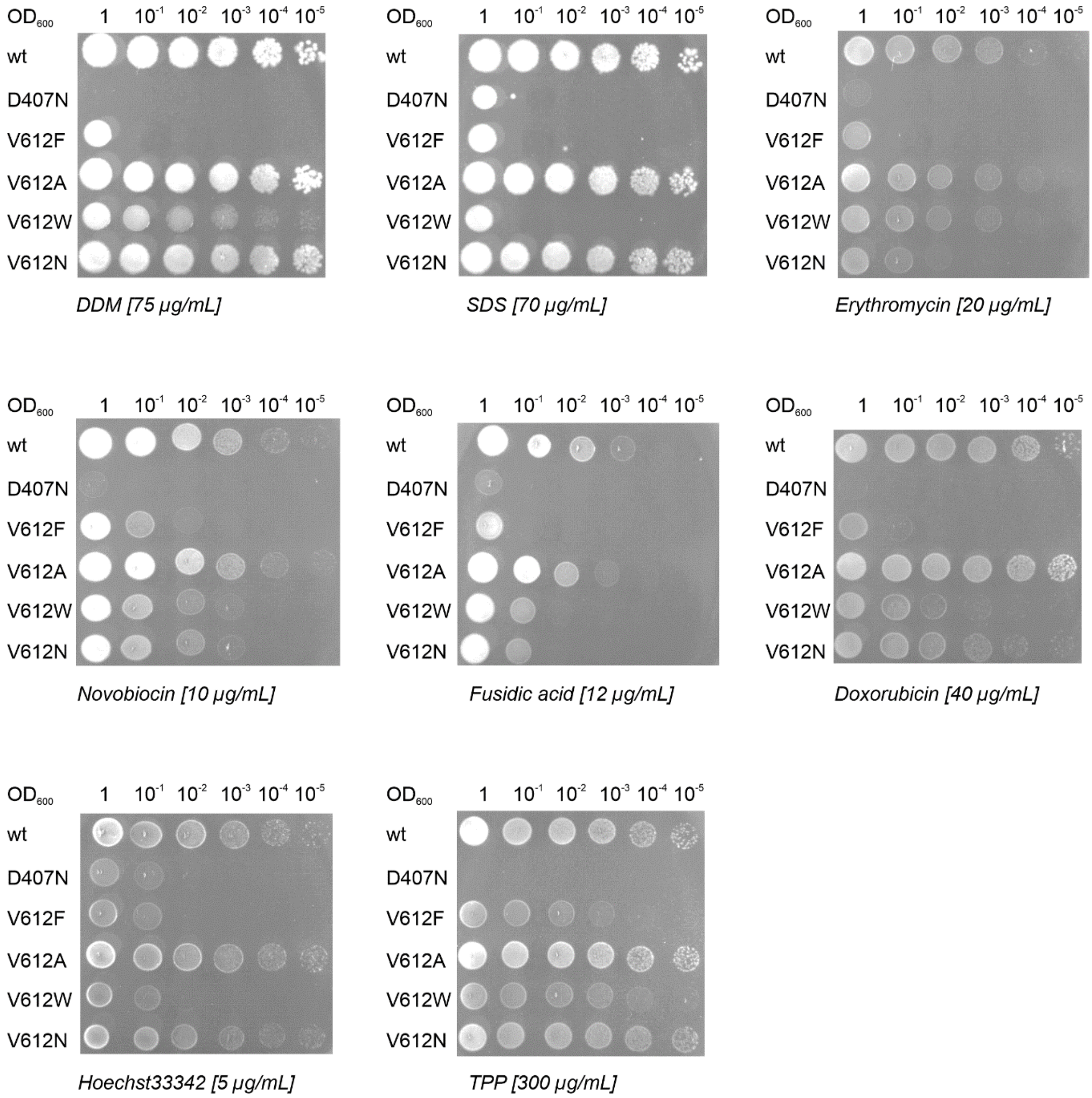
OD600 1 10-1 10-2 10-3 10-4 10-5
wt
D407N
V612F
V612A
V612W
V612N
DDM [75 µg/mL]
SDS [70 µg/mL]
Erythromycin [20 µg/mL]
Novobiocin [10 µg/mL]
Fusidic acid [12 µg/mL]
Doxorubicin [40 µg/mL]
Hoechst33342 [5 µg/mL]
TPP [300 µg/mL]

Biological replicate 1, technical replicate 2

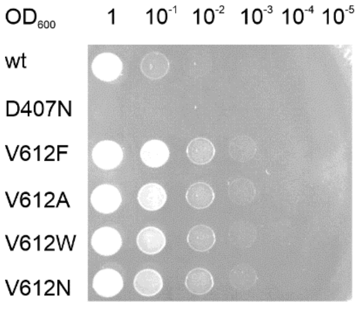


*Chloramphenicol [1.5 µg/mL]*

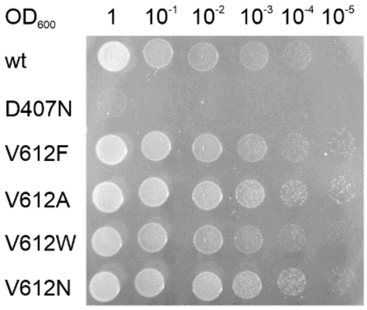


*Thiamphenicol [12 µg/mL]*

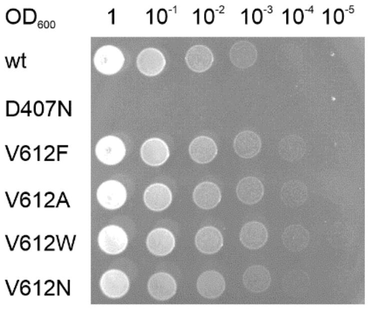


*Linezolid [20 µg/mL]*

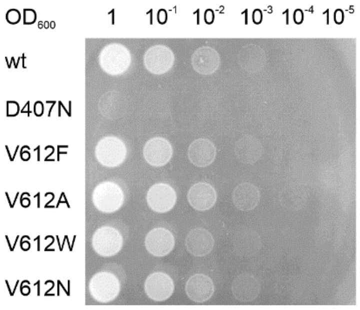


*Tetracycline [0.3 µg/mL]*

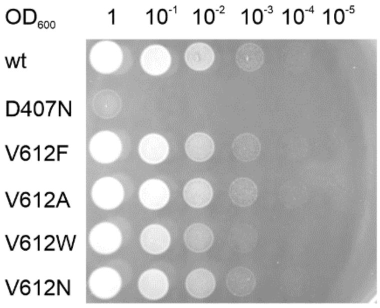


*Oxytetracycline [0.25 µg/mL]*

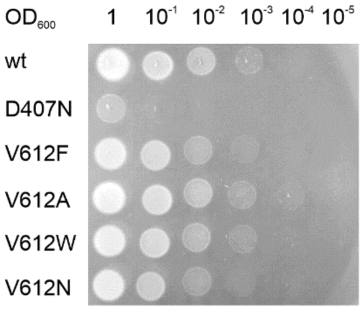


*Tigecycline [0.125 µg/mL]*

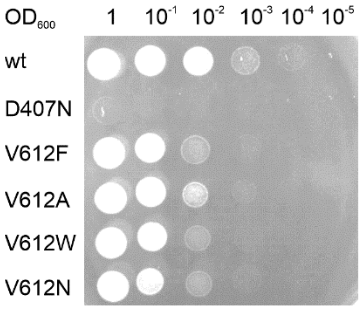


*Chlortetracycline [0.2 µg/mL]*

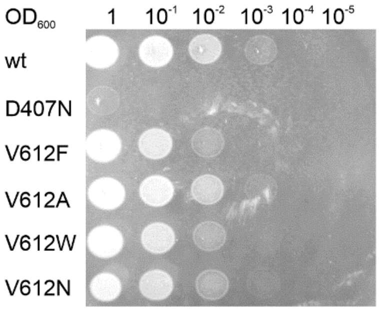


*Doxycycline [0.5 µg/mL]*

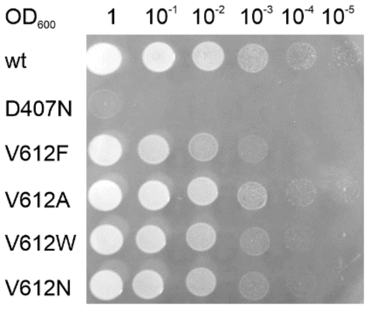


*Minocycline [0.3 µg/mL]*

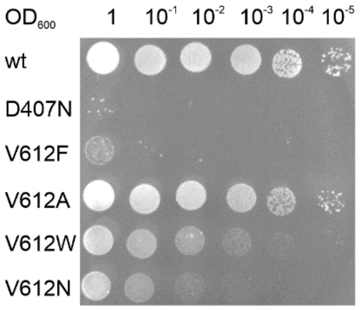


*Dicloxacillin [36 µg/mL]*

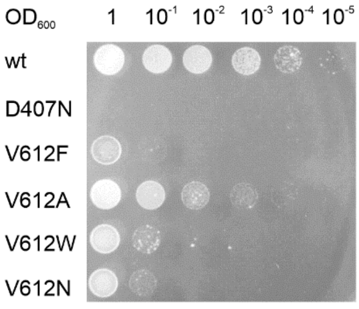


*Oxacillin [16 µg/mL]*

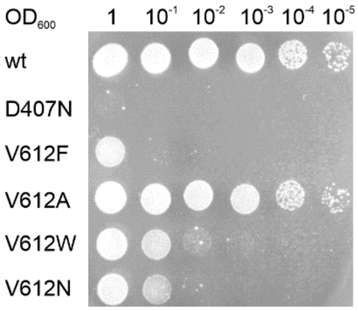


*Piperacillin [0.075 µg/mL]*

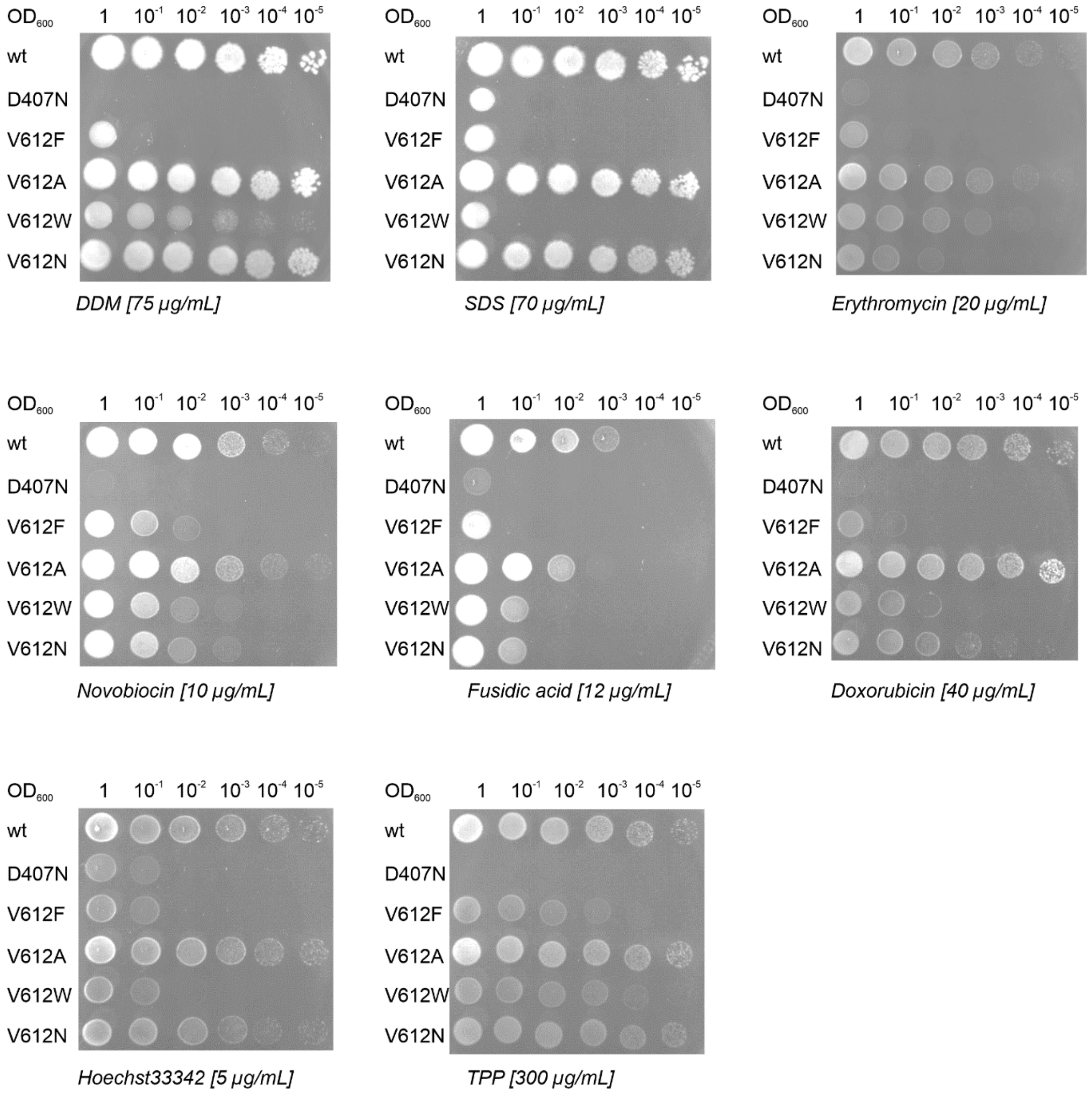
OD600
1
10-1
10-2
10-3
10-4
10-5
wt
D407N
V612F
V612A
V612W
V612N
DDM [75 µg/mL]
SDS [70 µg/mL]
Erythromycin [20 µg/mL]
Novobiocin [10 µg/mL]
Fusidic acid [12 µg/mL]
Doxorubicin [40 µg/mL]
Hoechst33342 [5 µg/mL]
TPP [300 µg/mL]

Biological replicate 2, technical replicate 1

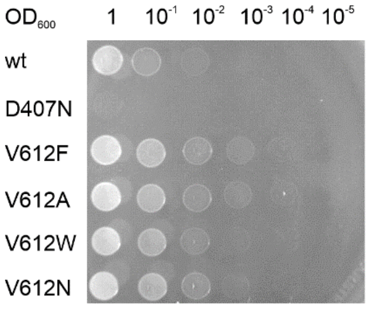

*Chloramphenicol [1.5 µg/mL]*

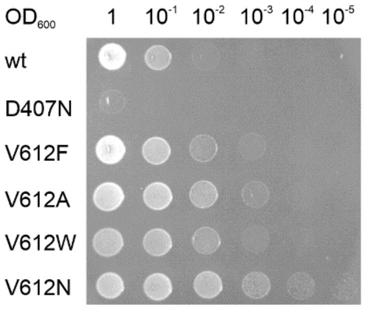

*Thiamphenicol [12 µg/mL]*

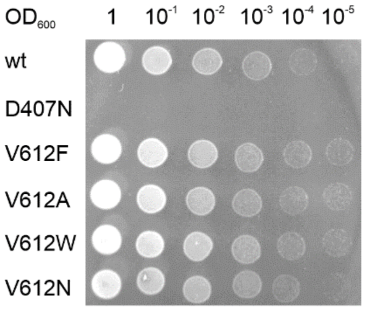

*Linezolid [20 µg/mL]*

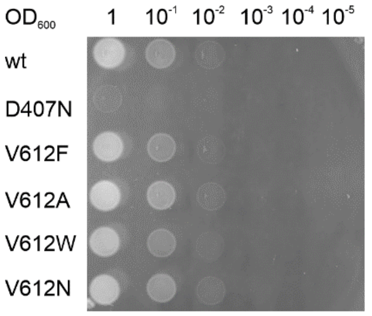

*Tetracycline [0.3 µg/mL]*

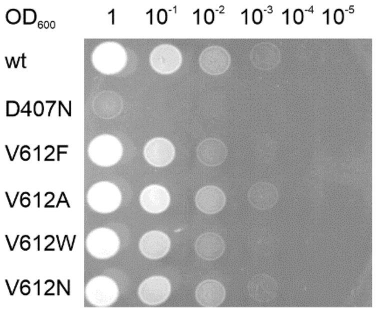

*Oxytetracycline [0.25 µg/mL]*

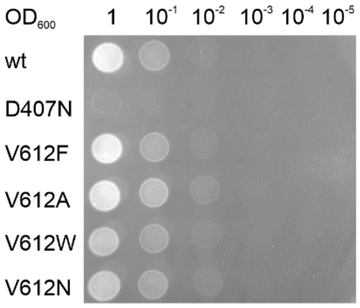

*Tigecycline [0.125 µg/mL]*

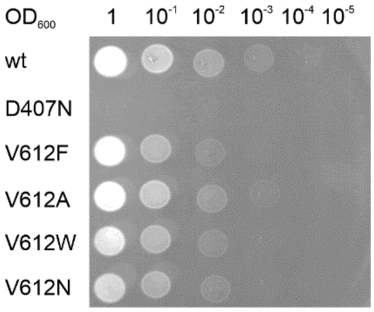

*Chlortetracycline [0.2 µg/mL]*

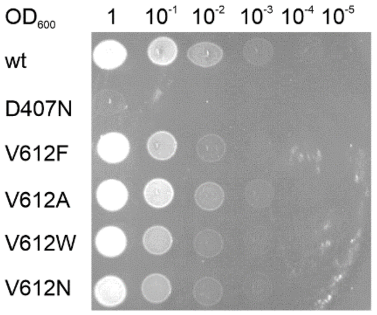

*Doxycycline [0.5 µg/mL]*

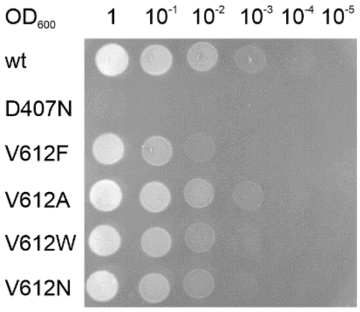

*Minocycline [0.3 µg/mL]*

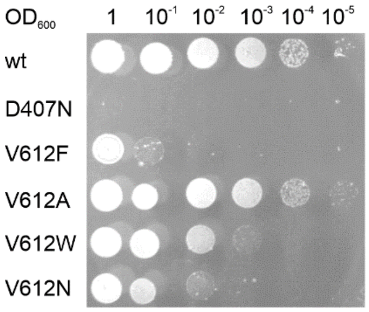

*Dicloxacillin [36 µg/mL]*

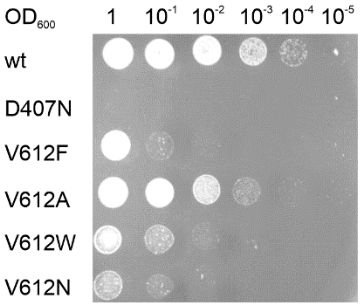

*Oxacillin [16 µg/mL]*

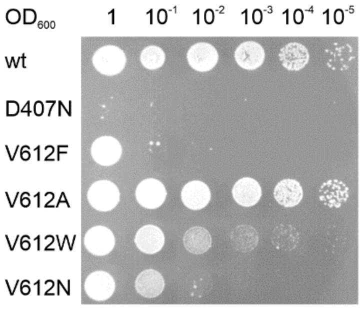

*Piperacillin [0.075 µg/mL]*

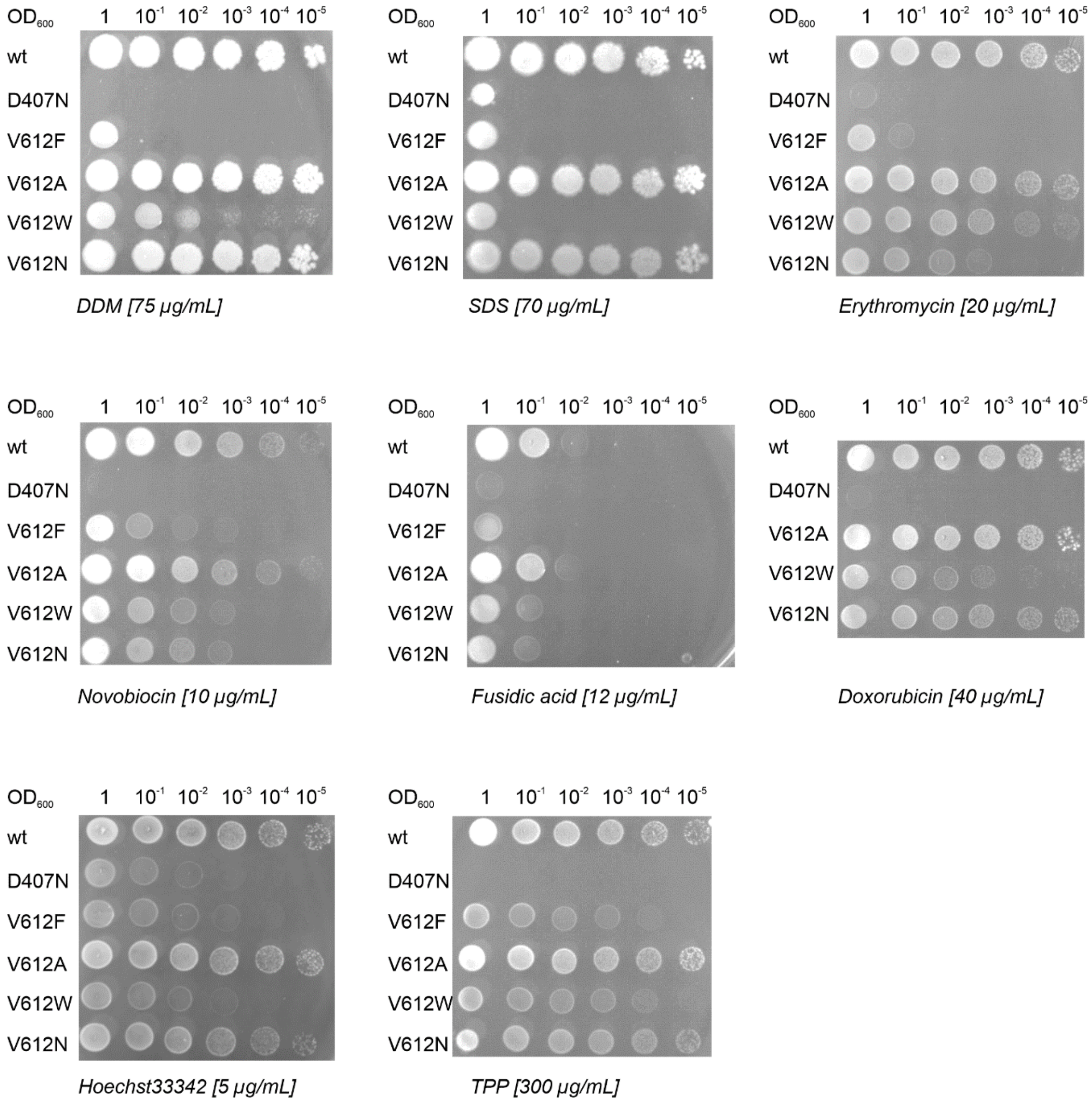

OD600 1 10-1 10-2 10-3 10-4 10-5
wt
D407N
V612F
V612A
V612W
V612N
DDM [75 µg/mL]
SDS [70 µg/mL]
Erythromycin [20 µg/mL]
Novobiocin [10 µg/mL]
Fusidic acid [12 µg/mL]
Doxorubicin [40 µg/mL]
Hoechst33342 [5 µg/mL]
TPP [300 µg/mL]

Biological replicate 2, technical replicate 2

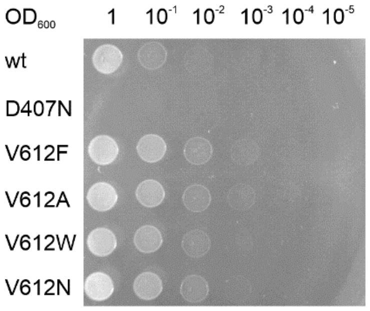


*Chloramphenicol [1.5 µg/mL]*

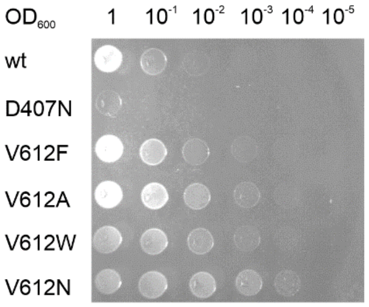


*Thiamphenicol [12 µg/mL]*

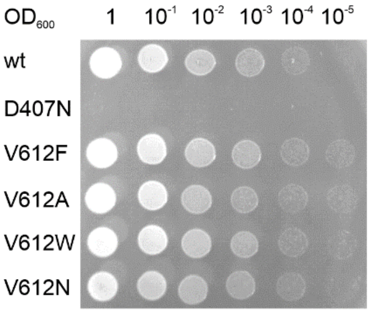


*Linezolid [20 µg/mL]*

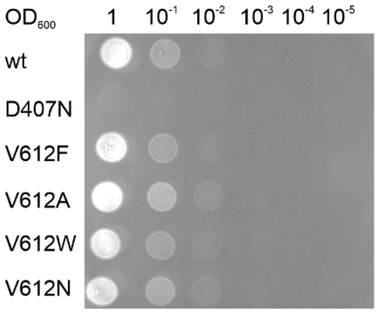


*Tetracycline [0.3 µg/mL]*

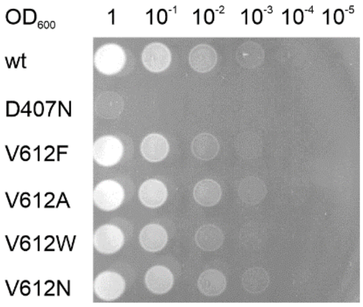


*Oxytetracycline [0.25 µg/mL]*

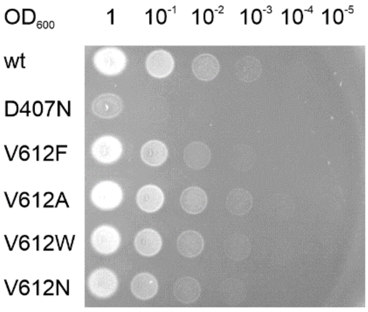


*Tigecycline [0.125 µg/mL]*

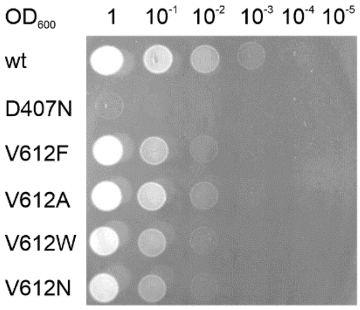


*Chlortetracycline [0.2 µg/mL]*

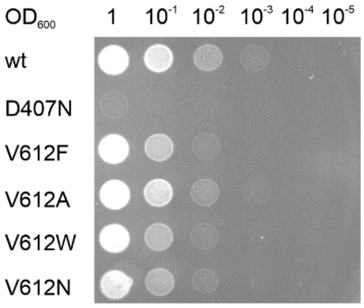


*Doxycycline [0.5 µg/mL]*

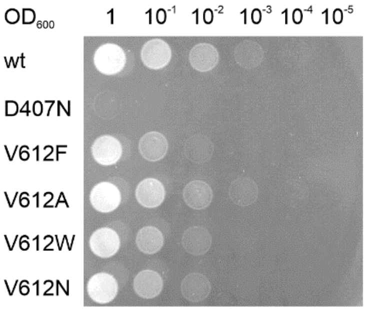


*Minocycline [0.3 µg/mL]*

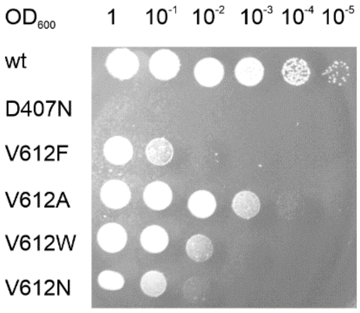


*Dicloxacillin [36 µg/mL]*

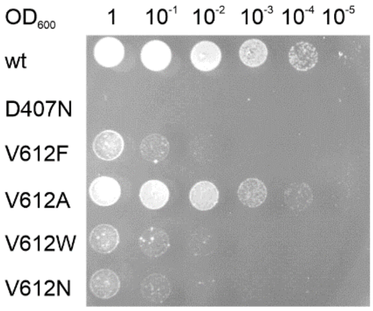


*Oxacillin [16 µg/mL]*

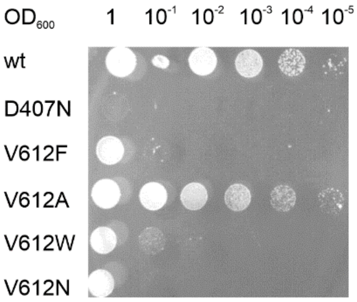


*Piperacillin [0.075 µg/mL]*

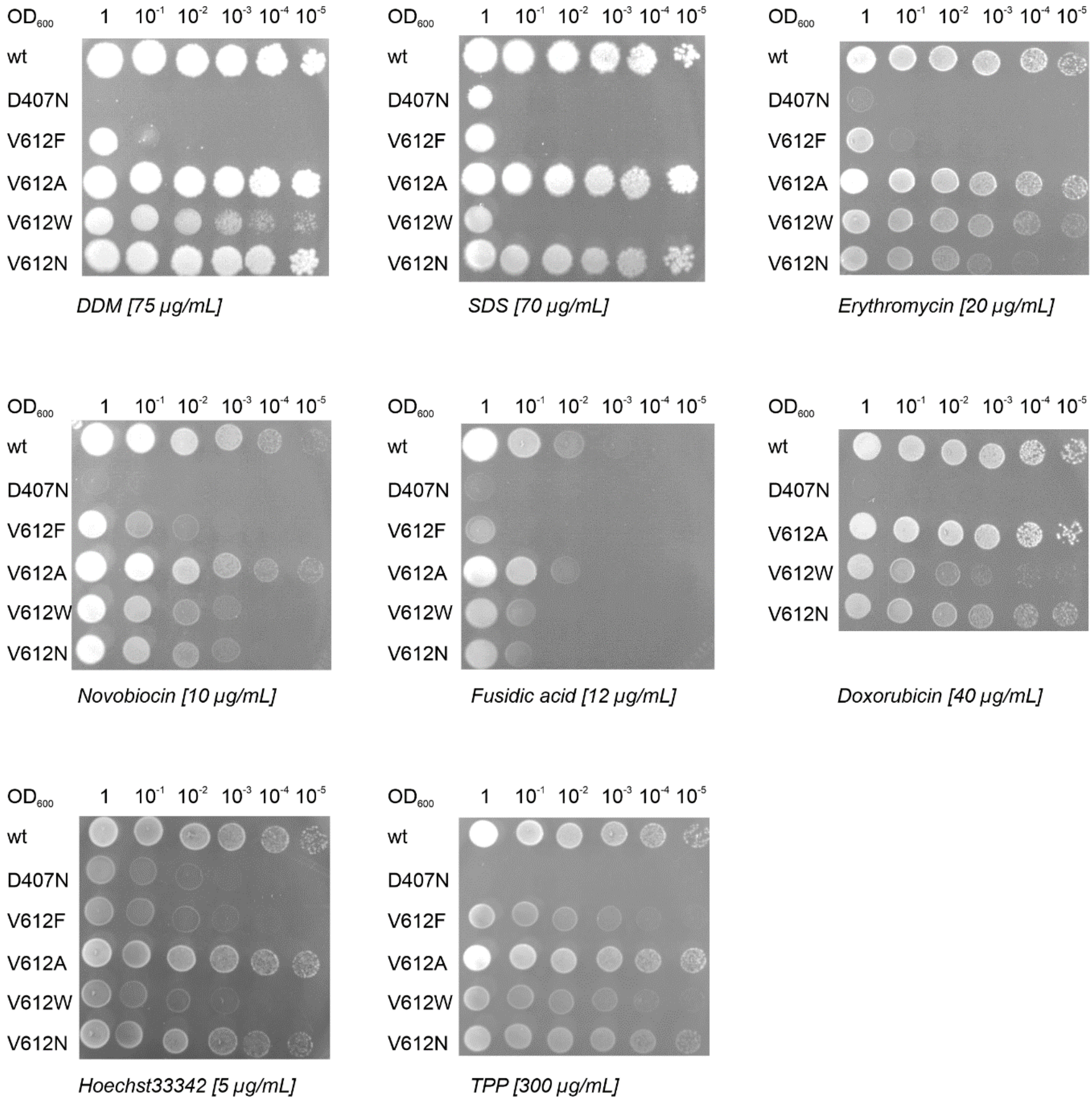
OD600
1
10-1
10-2
10-3
10-4
10-5
wt
D407N
V612F
V612A
V612W
V612N
DDM [75 µg/mL]
SDS [70 µg/mL]
Erythromycin [20 µg/mL]
Novobiocin [10 µg/mL]
Fusidic acid [12 µg/mL]
Doxorubicin [40 µg/mL]
Hoechst33342 [5 µg/mL]
TPP [300 µg/mL]

Biological replicate 3, technical replicate 1

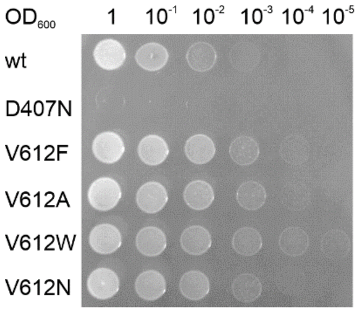


*Chloramphenicol [1.5 μg/mL]*

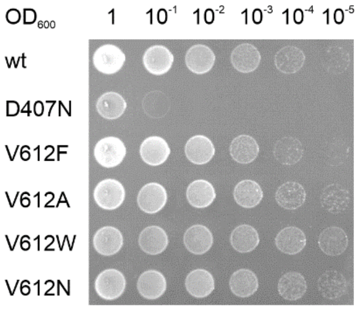


*Thiamphenicol [12 μg/mL]*

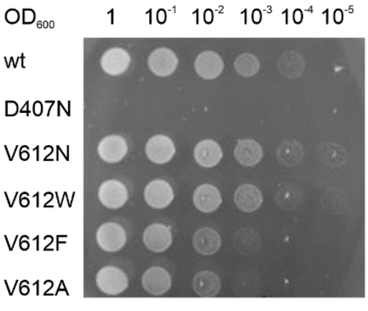


*Linezolid [20 μg/mL]*

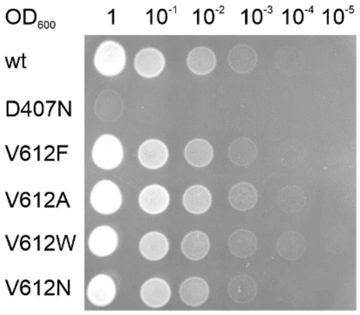


*Tetracycline [0.3 μg/mL]*

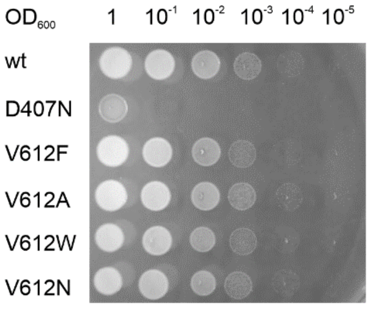


*Oxytetracycline [0.25 μg/mL]*

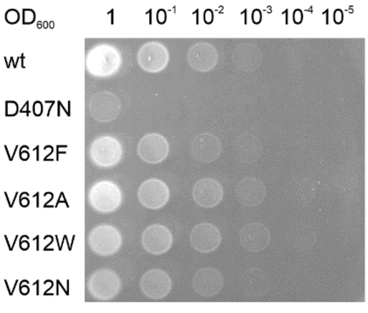


*Tigecycline [0.125 μg/mL]*

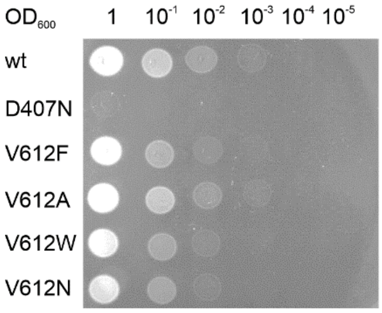


*Chlortetracycline [0.2 μg/mL]*

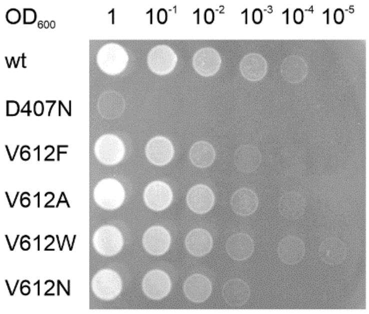


*Doxycycline [0.5 μg/mL]*

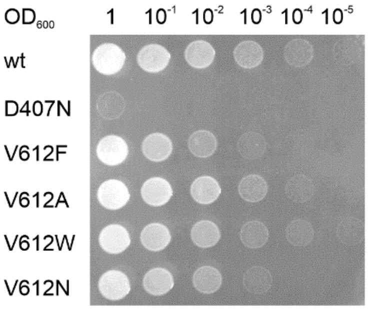


*Minocycline [0.3 μg/mL]*

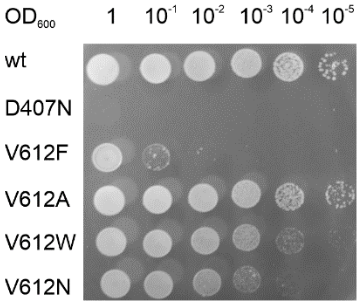


*Dicloxacillin [36 μg/mL]*

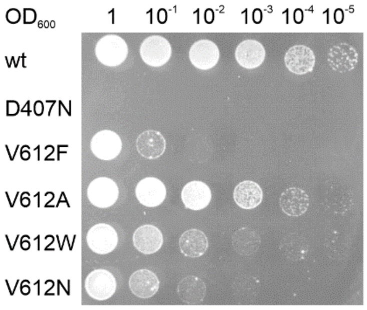


*Oxacillin [16 μg/mL]*

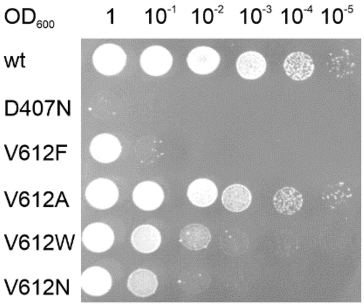


*Piperacillin [0.075 μg/mL]*

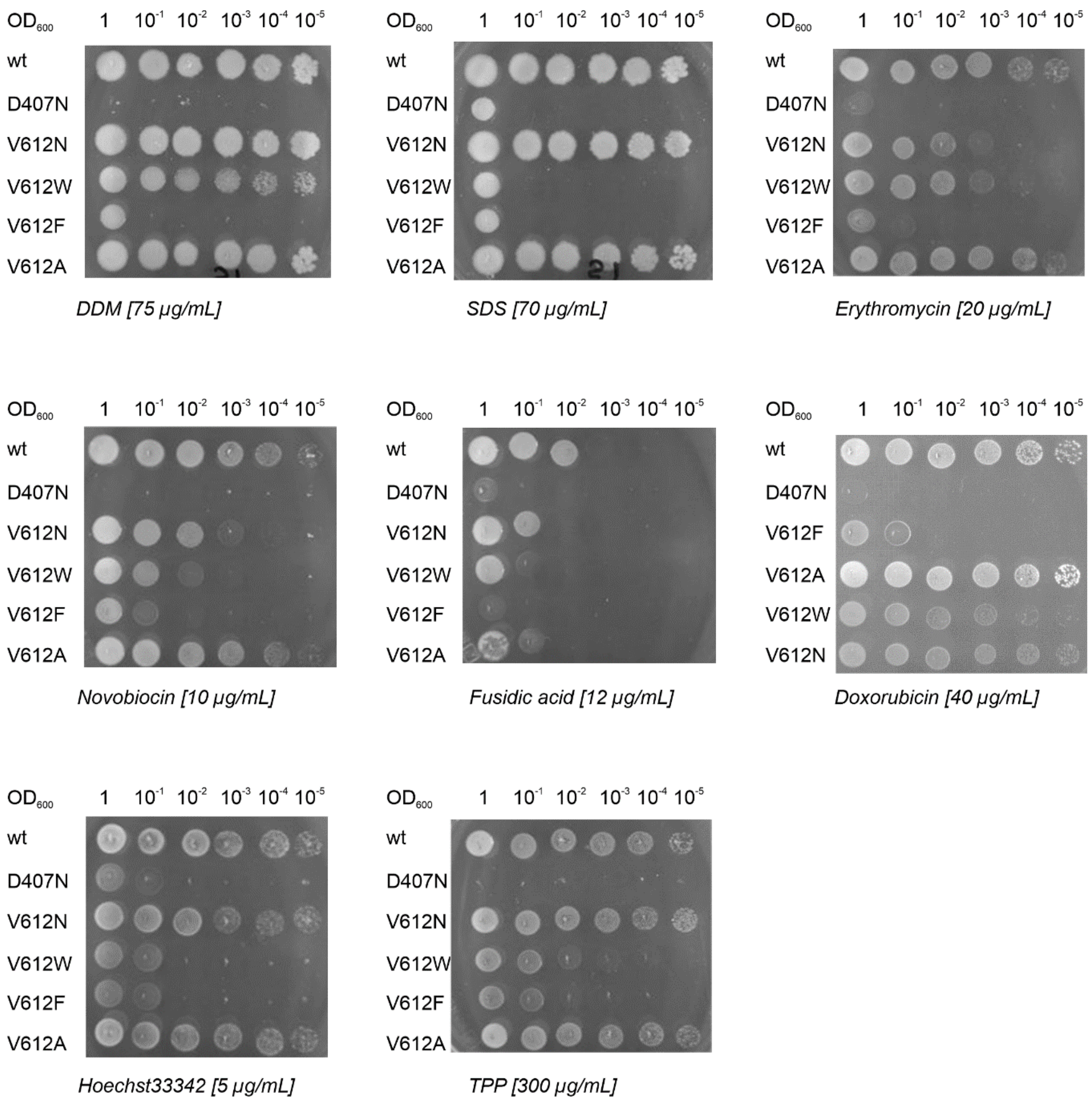

OD600 1 10-1 10-2 10-3 10-4 10-5
wt
D407N
V612N
V612W
V612F
V612A
DDM [75 µg/mL]
OD600 1 10-1 10-2 10-3 10-4 10-5
wt
D407N
V612N
V612W
V612F
V612A
SDS [70 µg/mL]
OD600 1 10-1 10-2 10-3 10-4 10-5
wt
D407N
V612N
V612W
V612F
V612A
Erythromycin [20 µg/mL]
OD600 1 10-1 10-2 10-3 10-4 10-5
wt
D407N
V612N
V612W
V612F
V612A
Novobiocin [10 µg/mL]
OD600 1 10-1 10-2 10-3 10-4 10-5
wt
D407N
V612N
V612W
V612F
V612A
Fusidic acid [12 µg/mL]
OD600 1 10-1 10-2 10-3 10-4 10-5
wt
D407N
V612F
V612A
V612W
V612N
Doxorubicin [40 µg/mL]
OD600 1 10-1 10-2 10-3 10-4 10-5
wt
D407N
V612N
V612W
V612F
V612A
Hoechst33342 [5 µg/mL]
OD600 1 10-1 10-2 10-3 10-4 10-5
wt
D407N
V612N
V612W
V612F
V612A
TPP [300 µg/mL]

Biological replicate 3, technical replicate 2

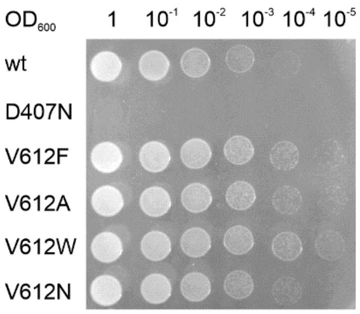


*Chloramphenicol [1.5 µg/mL]*

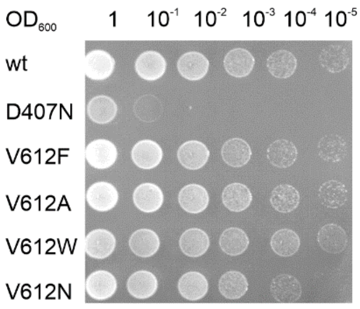


*Thiamphenicol [12 µg/mL]*

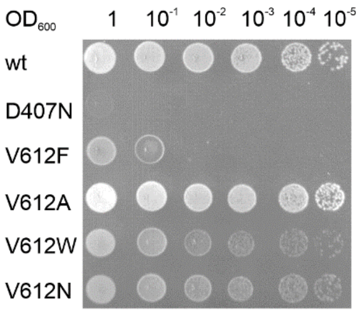


*Doxorubicin [40 µg/mL]*

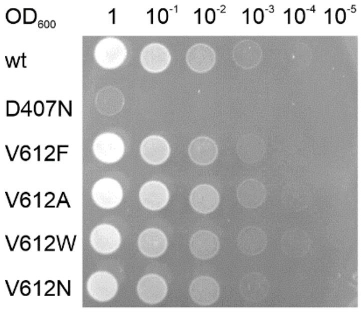


*Tetracycline [0.3 µg/mL]*

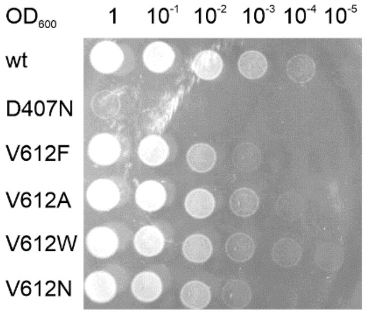


*Doxycycline [0.5 µg/mL]*

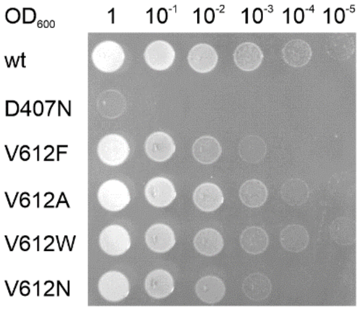


*Minocycline [0.3 µg/mL]*

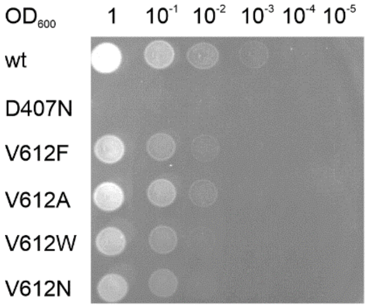


*Chlortetracycline [0.2 µg/mL]*

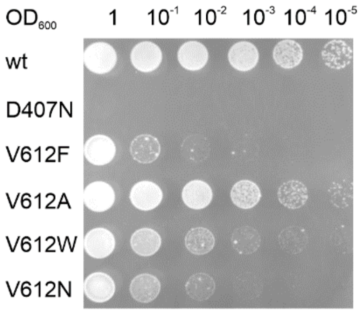


*Oxacillin [16 µg/mL]*
